# NOTCH1 Acts as a Tumor Suppressor That Induces Early Differentiation in Head and Neck Cancer

**DOI:** 10.1101/2025.04.25.650710

**Authors:** Chenfei Huang, Shhyam Moorthy, Qiuli Li, Kazi Mokim. Ahmed, Defeng Deng, Jiping Wang, Xiayu Rao, Jiexin Zhang, Yuanxin Xi, Jing Wang, Zhiyi Liu, Noriaki Tanaka, David A. Wheeler, Eve Shibrot, Rami Saade, Curtis R. Pickering, Tong-Xin Xie, Adel K. El-Naggar, Abdullah A. Osman, Kunal Rai, Patrick A. Zweidler-McKay, John V. Heymach, Lauren A. Byers, Faye M. Johnson, Vlad C. Sandulache, Jeffrey N. Myers, Pedram Yadollahi, Mitchell J. Frederick

## Abstract

We identified frequent inactivating notch1 mutations in HNSCC over a decade ago, indicating its role as a tumor suppressor—unlike its oncogenic function in leukemias and salivary gland tumors. However, there has been much debate in the literature regarding a possible oncogenic role in HNSCC as well, based on reports that notch1 signaling drives tumor growth and a cancer stem cell phenotype in some HNSCC tumor lines and patient samples. Clarifying whether NOTCH1 occasionally functions as an oncogenic driver in HNSCC is crucial to the prognosis and personalized therapy of patients with either wild-type or mutated NOTCH1. Here we present a systematic and comprehensive investigation unequivocally demonstrating that notch1 signaling functions as a tumor suppressor in HNSCC regardless of mutation or activation status and leads to reduction in frequency of cancer stem cells. We develop a robust gene expression signature of notch1 activation based on experimental data that when applied to patient samples shows notch1 signaling is associated with very early differentiation, an altered tumor microenvironment, and better prognosis— consistent with a tumor suppressive role. Our work unifies the field by reconciling conflicting data and providing critical insights into the biological and clinical significance of the NOTCH1 pathway in HNSCC.

## INTRODUCTION

The genomic landscape of head and neck squamous cell carcinoma (HNSCC) is predominated by tumor suppressors (1–4), posing challenges for the development of molecular targeted therapies. Our team was among the first to identify frequent inactivating *NOTCH1* mutations in HNSCC (1,3) and aggressive cutaneous SCC (cSCC) (5), indicating a potential tumor suppressive role. Subsequent investigations have confirmed similar mutation patterns in HNSCC (4,6,7), cSCC (8), and SCCs of the lung (LUSC) (9) and esophagus (ESCC) (10) through The Cancer Genome Atlas (TCGA) and independent studies, solidifying *NOTCH1* as one of the most commonly mutated genes across various SCCs. The presence of mutations in extracellular ligand binding domains and truncating mutations throughout the *NOTCH1* gene in SCCs aligns with its proposed tumor suppressor function (11), supported by earlier mouse studies demonstrating increased skin tumors upon conditional *NOTCH1* knockout (12). Conversely, *NOTCH1* is an oncogenic driver in T cell acute lymphoblastic leukemia (T-ALL), where activating missense mutations cluster in the heterodimerization (HD) domain and truncating mutations occur in the C-terminal PEST sequence, leading to increased *NOTCH1* activation (11,13). This oncogenic role has also been reported in adenoid cystic carcinomas originating from the salivary gland (14), highlighting NOTCH1’s context-specific dual role in cancer biology.

Although *NOTCH1* plays opposing roles in cancers originating from different tissues, multiple studies suggest that it may also have a dual function within HNSCC tumors of the same histology, including an oncogenic role (7,15–19). *NOTCH1* mutations found in the HD and Abruptex region presumed to be activating have been reported for several cohorts of Asian HNSCC patients (20,21), although subsequent cloning of an Abruptex mutation later revealed that it was inactivating (22). Nevertheless, studies have correlated overexpression of *NOTCH1* RNA and protein with poor prognosis in HNSCC (15,17,18,23), and utilized pharmacological inhibition of NOTCH1 signaling or gene knockdown to link NOTCH1 activation to proliferation (16,19,23), *in vivo* tumor growth (18,24), tumor spheroid formation (19,24,25), and resistance to chemotherapy (25). In contrast, we previously demonstrated that ectopic expression of activated intracellular NOTCH1 (ICN1) inhibits proliferation and *in vivo* tumor growth in multiple *NOTCH1*-mutant HNSCC cell lines (2). The latter is supported by a report that activated NOTCH1 detected by immunohistochemistry (IHC) in the tumor nuclei correlated with better survival in HNSCC patients (26).

Clarifying whether the NOTCH1 pathway occasionally functions as an oncogenic driver in HNSCC is crucial not only for academic discourse but also for the prognosis and personalized therapy of SCC patients with either wild-type (WT) or mutated *NOTCH1*. While NOTCH1 inhibition has been proposed for HNSCC tumors where the pathway is deemed oncogenic, we reported that HNSCC cell lines harboring inactivating *NOTCH1* mutations are highly sensitive to PI3K inhibitors (27–29), underscoring the complex interplay of signaling pathways in SCC. Here, we present a systematic and comprehensive analysis demonstrating that *NOTCH1* functions as a tumor suppressor in HNSCC regardless of mutation or activation status, despite its potential to induce pseudo-stem cell-like properties *in vitro.* We further demonstrate that restoration or activation of NOTCH1 signaling induces a program of very early differentiation―also manifested in human HNSCC primary tumors― that reshapes the tumor microenvironment and may influence which tumor dependencies can be clinically targeted successfully. Our findings challenge prevailing models that consider the de-differentiated state of squamous cell carcinomas to be irreversible due to genetic mutations, offering deeper insights into the mechanisms of tumor plasticity.

## RESULTS

### NOTCH1 Restoration Inhibits Growth in Two-Dimensional Cultures and Alters Morphology of Head and Neck Squamous Cell Carcinoma (HNSCC) Cell Lines Harboring Inactivating *NOTCH1* Mutations

To understand the consequences of NOTCH1 signaling in HNSCC lines harboring *NOTCH1* inactivating mutations (Table S1), we re-expressed WT full-length *NOTCH1* (NFL1) receptors using a bicistronic retroviral vector harboring an IRES-EGFP tag. This allowed purification and testing of NFL1 expressing cells while avoiding artifacts of long-term selection. Following NFL1 overexpression, cells were continuously cultured (1-10 days) on plates pre-coated with either recombinant NOTCH1-ligand Jagged1 fused to an FC-receptor (JAG1) or immobilized control FC protein. We detected cleaved/activated intracellular NOTCH1 (cl-NOTCH1) protein only in *NOTCH1* mutants infected with NFL1 and not empty vector control virus (MigR1), which increased substantially after just 16 h of growth on JAG1 compared to FC control protein (Fig. S1A) and confirmed that endogenous *NOTCH1* mutations were indeed inactivating. NFL1 overexpression alone, without external JAG1, modestly decreased colony formation in both UMSCC47 and UMSCC22A (Fig. S1B and S1C) over a 10-day period. However, when grown in the presence of immobilized JAG1 ligand, NFL1 expression led to a significant reduction in colony formation in four different *NOTCH1*-mutant cell lines when compared to growth on control FC protein (Fig. S1B and S1C).

Reduced colony growth in NOTCH1-mutant UMSCC22A expressing WT NFL1 that were exposed to JAG1 was accompanied by the onset of profound morphological changes after 3 to 5 days, which included a vast reduction in cell size and compact growth as loosely attached spheroids (Fig. S1D). A fraction of NOTCH1-mutant HN4 and HN31 cells expressing NFL1 became spindle-shaped after 3 to 5 days growth on JAG1 and positive for the senescent marker beta-galactosidase (β-Gal) (Fig. S2A and S2B). Likewise, the loosely formed spheroids produced by NOTCH1-mutant UMSCC22A cells that express NFL1 and were grown on JAG1 showed considerable β-Gal staining (Fig. S2A) Collectively, reactivating NOTCH1 signaling in mutant tumors profoundly inhibited cell growth in two-dimensional cultures and led to altered morphology and senescence.

### HNSCC Cell Lines Harboring Wild Type *NOTCH1* Show Similar Patterns of Growth Inhibition and Altered Morphology Following NOTCH1 Activation

Next, we examined the morphological and proliferation phenotypes associated with NOTCH1 pathway activation in four randomly selected HNSCC cell lines expressing endogenous WT *NOTCH1* receptors (Table S1). Expression of full-length transmembrane NOTCH1 (Tm-NOTCH1) receptor proteins at various levels was confirmed by Western blotting (Fig. 1A). Levels of cl-NOTCH1 protein were examined in three of the cell lines grown on control FC protein and found to be barely detectable but increased following a brief 16 h exposure to JAG1 (Fig. 1A). Extended cultivation of *NOTCH1* WT tumors on immobilized JAG1 inhibited colony formation (Fig. 1B) in four cell lines tested (PJ34, 183, CAL27, and UMSCC1), which was really striking in 3 out of the 4 cell lines where ligand-induced NOTCH1 activation had been confirmed (e.g. Fig. 1A). The growth inhibition was consistent with that observed for *NOTCH1* mutants after restoring NOTCH1 signaling. In the fourth NOTCH1 WT cell line, UMSCC1, JAG1 induced mild but significant reduction in colony formation (Fig. 1B). Remarkably, growth of 183 and PJ34 cell lines on JAG1 (but not control FC protein) led to the very same morphological transformation found earlier in UMSCC22A, characterized by substantial cell shrinkage and the formation of loosely attached compacted tumor spheroids (Fig. 1C). These spheroids also displayed positive staining for β-Gal (Fig. 1D).

**Figure 1.**
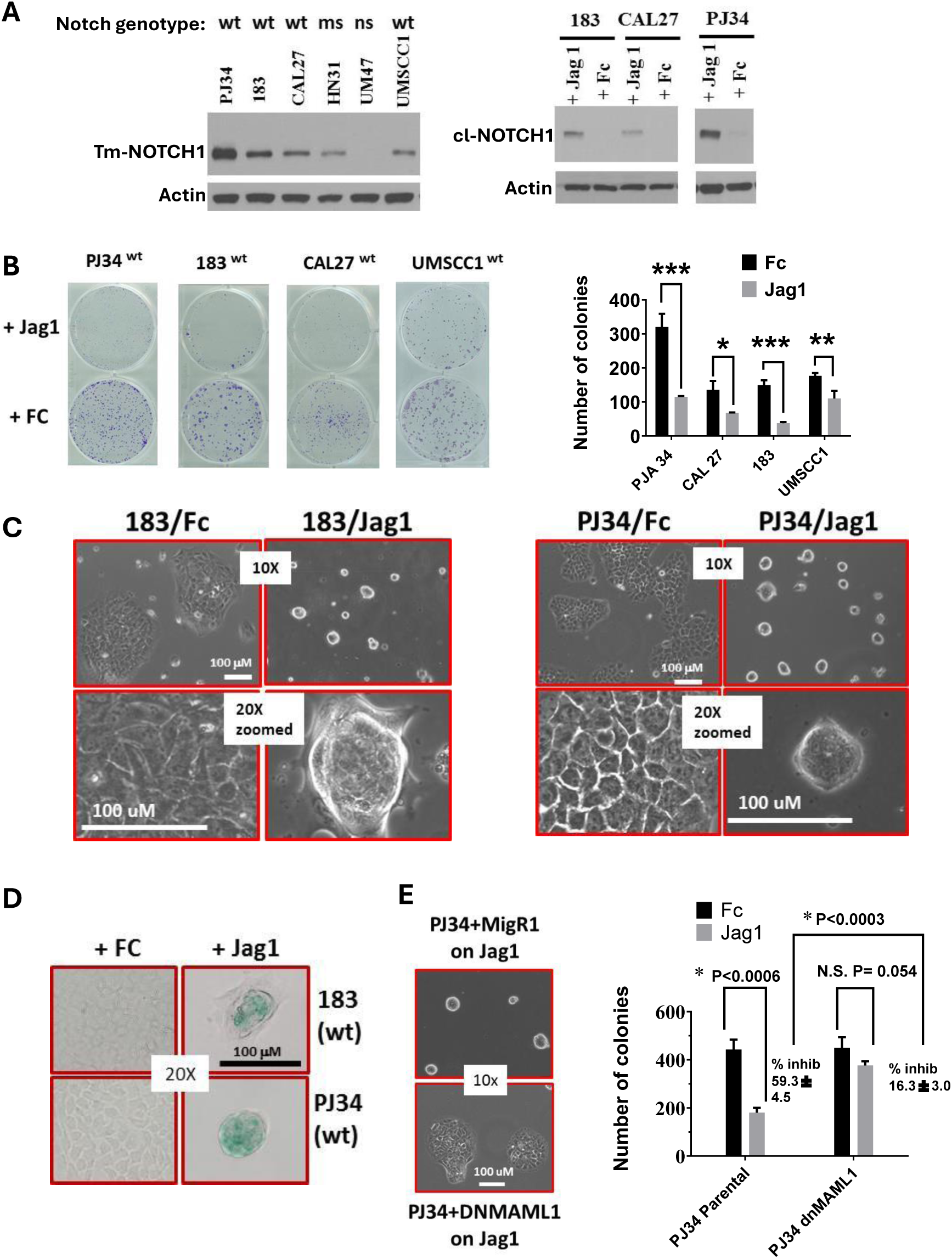
NOTCH signaling alters cell morphology and inhibits growth in two-dimensional cultures. **A.** Expression of full length transmembrane NOTCH1 (Tm-NOTCH1) in cells with WT *NOTCH1* (PJ34, 183, CAL27, and UMSCC1) and cleaved activated NOTCH1 (cl-NOTCH1) protein after 16 h growth on immobilized Notch1 ligand (JAG1) or FC control protein (FC). **B.** Extended growth (e.g. 8-12 days) on JAG1 ligand significantly reduced colony formation compared to control FC protein in four *NOTCH1* WT cell lines. **C.** Growth on JAG1, but not control FC induces morphological transformation of cell lines with WT *NOTCH1* (183 and PJ34) observed by 5 days, characterized by reduced cell size and formation of loosely attached tumor spheroids. **D.** Tumor spheroids induced by growth on JAG1 express the senescence marker b-gal. **E.** Ectopic expression of dominant negative MAML1, which inhibits NOTCH1 mediated transcriptional regulation, prevents JAG1-induced tumor spheroid formation and reverses inhibition of colon formation in PJ34 cells.

The growth inhibition and associated morphological changes induced by JAG1 exposure in PJ34 were effectively reversed by expressing a dominant negative form of Mastermind-like 1 (Fig. 1E), known to inhibit NOTCH family signaling. Intrigued, we aimed to dissect the individual phenotypic contributions of NOTCH1 and NOTCH2 signaling in PJ34, since both receptors can be triggered by their shared ligand JAG1. Through CRISPR knockout (KO) experiments targeting *NOTCH1, NOTCH2*, or both genes (Supplementary Fig. S3A), we observed that knockout of either NOTCH gene only partially rescued PJ34 from the growth inhibition induced by JAG1 (Supplementary Fig. S3B). However, double knockout of both genes (N1N2KO) almost completely reversed the JAG1-mediated growth inhibition (Supplementary Fig. S3B) and prevented morphological formation of tumor spheroids (Supplementary Fig. S3E). Re-expression of NFL1 alone in N1N2KO cells was sufficient to restore JAG1-induced NOTCH1 activation (Supplementary Fig. S3C), growth inhibition (Supplementary Fig. S3D), and morphological changes (Supplementary Fig. S3E).

### NOTCH1 Activation Does not Drive Proliferation in HNSCC Tumors with High Endogenous NOTCH1 Signaling

Because previous studies linked NOTCH1 signaling to proliferation and cancer stem cell like behavior in HNSCC cell lines, we investigated the function of NOTCH1 signaling in tumors with high endogenous levels of activated cl-NOTCH1 initially identified by reverse phase protein arrays (RPPA). Levels of cl-NOTCH1 along with 155 other proteins/phosphoproteins, including total NOTCH1 were measured across 53 different HNSCC cell lines with known *NOTCH1* and *NOTCH2* mutational status (Table S1). As predicted, RPPA levels of cl-NOTCH1 were significantly lower in cell lines harboring *NOTCH1* mutations (P =0.03, Fig. S4A). Western blotting (Supplementary Fig. S4B and S4C) confirmed relatively high levels of baseline cl-NOTCH1 protein in six cell lines (FaDu, SCC61, SCC15, PCI24, MDA1986LN, and MDA686LN) identified by RPPA, compared to two of the *NOTCH1* WT cell lines, PJ34 and CAL27, utilized earlier and found to undergo JAG1-mediated growth inhibition (e.g. Fig. 1B, and Fig. S4B). To evaluate the necessity of NOTCH1 signaling in these cells with high cl-NOTCH1 levels, we inhibited the formation of cl-NOTCH1 using 250 or 500 nM of the gamma secretase inhibitor DBZ in all six cell lines (Fig. 2A and Supplementary Fig. S4D). The inhibition of NOTCH1 activation persisted 48 to 72 hours post-DBZ treatment and continuous exposure to DBZ (refreshed every 48 hours) failed to inhibit clonogenic growth in NOTCH1-wild type cells FADU, PCI24, SCC61, SCC15, or MDA686LN (Fig. 2B and 2C). On the contrary, there was even a slight increase in colony formation for two of the cell lines with DBZ. Although MDA1986 LN failed to form colonies in the absence of treatment, the growth of MDA1986 LN was unaffected by DBZ in standard proliferation assays (not shown).

**Figure 2.**
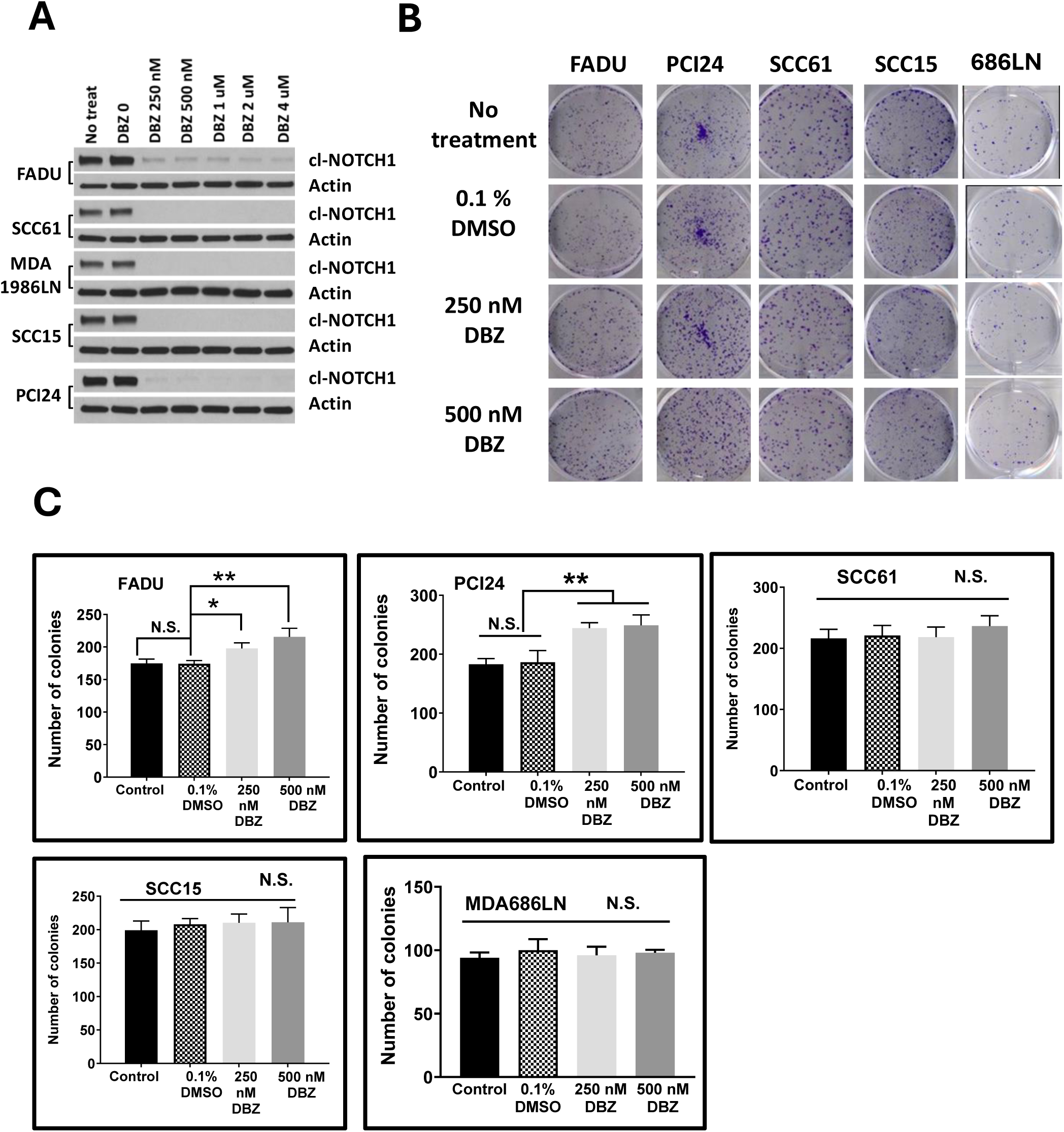
NOTCH1 is not a driver of cell growth in multiple HNSCC cell lines with high endogenous NOTCH1 activation. **A.** Western blot validation of high basal levels of cl-NOTCH1protein in 5 NOTCH1 WT HNSCC cell lines with different genomic backgrounds and persistent inhibition of NOTCH1 signaling after 72 h treatment with various doses of the NOTCH1 inhibitor DBZ. **B**. Staining of colony formation in the presence or absence of continuous treatment with DBZ inhibitor (replaced every 48 h) for the duration of culture. **C.** Quantitation of colony formation shows no difference in growth after continued treatment with DBZ, except for FaDu cells which grew slightly better after inhibition of NOTCH1 signaling.

### Proteins Correlating with NOTCH1 activation in HNSCC cell lines

The top protein correlating with cl-NOTCH1 levels among analytes analyzed by RRPA was total NOTCH1 (r = 0.679, AdjP = 3.50E-06, Table S2), followed by FBXW7 (r= 0.612) and EZH2 (r = 0.608). FBXW7 is known to degrade active cl-NOTCH1 in the nucleus and likely represents negative feedback, while EZH2 is a histone methyltransferase that represses gene expression. NRF2, which is a stress-induced transcriptional activator responsive to reactive oxygen species (ROS) was also positively correlated (r =0.449), as was KEAP1 (0.331) that negatively regulates NRF2, and NF-kB-p65 (r = 0.382) which also responds to ROS. Among significantly anti-correlated proteins were the receptor tyrosine kinase AXL (r = −0.457) that regulates survival and proliferation and fibronectin (r =-0.385) an extracellular matrix protein that can mediate binding of fibroblasts.

### Persistent NOTCH1 signaling downregulates Proto-Oncogenes, Growth Factors, and Integrins While Increasing Expression of Early Differentiation Markers

To elucidate the molecular mechanisms and understand phenotypes associated with *prolonged activated NOTCH1 signaling*, we examined alterations in gene expression in *NOTCH1*-WT HNSCC cell lines (PJ34 and 183) induced by growth on JAG1 ligand for five days. A two-way ANOVA identified genes downregulated (Table S3) and upregulated (Table S4) due to growth on JAG1 as a main effect (e.g., JAG treatment) and for the individual cell lines PJ34 and 183 grown on JAG1 in post hoc analyses. We focused on the top significant genes that showed at least a 1.4-fold change in both cell lines and found 50 genes upregulated and 70 downregulated because of JAG1 exposure. Gene Ontology (GO) enrichment analysis of these 120 altered genes identified pathways related to proliferation, differentiation, cell adhesion, cytokine production, response to oxygen containing compounds, and cytokine production (Supplementary Table S5 and S6, and Supplementary Fig. S5). Among the top genes downregulated by NOTCH activation were three pro-survival/proliferation genes, *α-CATULIN* (*CTNNAL1*), *AXL*, and Epiregulin (*EREG*). Notably, *α-CATULIN* exhibited the most significant reduction in magnitude among all genes (e.g., 24-fold in 183) following NOTCH activation. *AXL* ranked among the top five genes with the greatest reduction in both cell lines and was consistent with the earlier finding of an inverse correlation between AXL protein and cl-NOTCH1 by RRPA (Supplementary Table S2). Multiple genes regulating cellular adhesion, including *ITGA3* (integrin α3), *ITGA5* (integrin α5), *LAMC2* (Laminin gamma 2), and *LAMC1*, were downregulated in both cell lines after NOTCH activation (Supplementary Table S3). Keratins 4 and 13 emerged as the top two upregulated genes in 183 cells (7.9- and 3.6-fold, respectively) and among the top 15 upregulated genes in PJ34 (2.0- and 1.8-fold, respectively) following NOTCH activation (Supplementary Table S4). Additionally, putative tumor suppressors such as *EPHA4, TP53INP1, PDCD4, DSP,* and *TXNIP* were among top genes upregulated by NOTCH activation. The specific pattern of integrins downregulated and keratins upregulated (Fig. 3A) mirrors what happens to normal oral mucosa during very early differentiation as basal stem cells divide and migrate upwards to the suprabasal layer of squamous epithelium. Collectively, the genes commonly regulated in both PJ34 and 183 cells after growth on ligand suggest a loss of proliferation and loss of substrate adhesion consistent with very early squamous cell differentiation.

**Figure 3.**
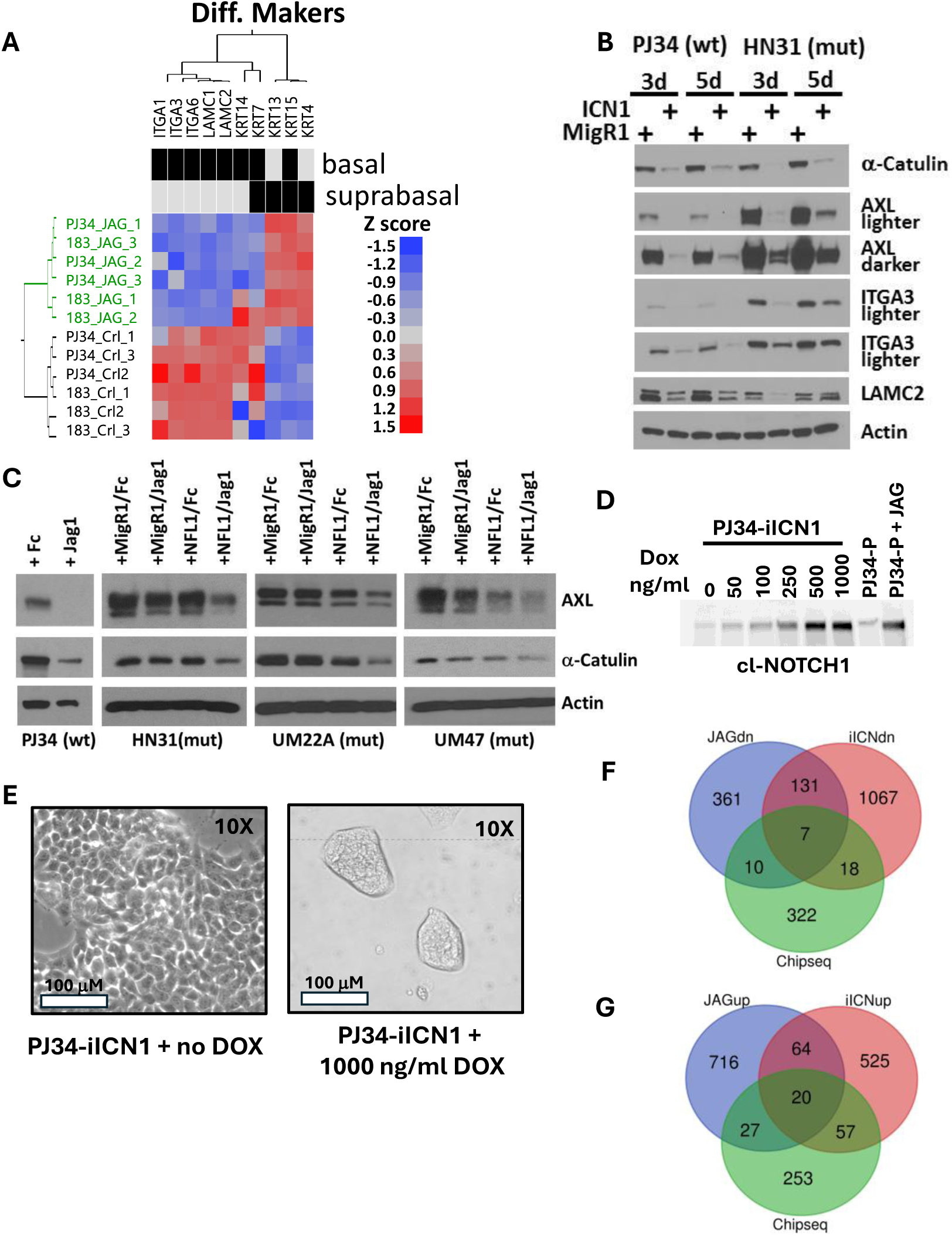
The activation of NOTCH1 signaling through multiple experimental approaches consistently downregulates oncogenic drivers and molecules mediating cell adhesion. **A.** Growth for 5 days on JAG1 significantly inhibited RNA expression of the basal cell marker genes ITGA1, ITGA3, ITGA6, LAMC1, and LAMC2 in two *NOTCH1* WT cell lines PJ34 and 183, while stimulating expression of the suprabasal maker genes KRT4and KRT13. **B.** Ectopic expression of cDNA encoding cl-NOTCH1decreased expression of ITGA3, LAMC2, AXL, and α-CATULIN at 3 days and 5-days post infection in a *NOTCH1* WT cell (PJ34) and in a *NOTCH1*-mutant cell (HN31). **C**. AXL and a-CATULIN protein levels decline in *NOTCH1* WT (PJ34) tumor cells grown on JAG1 (3 days) and in 3 different *NOTCH1* mutant cell lines (HN31, UMSCC22A, UMSCC47) when expression of WT full length NOTCH1 receptor (NFL1) is restored (NFL1) and then grown on JAG1 ligand for 3 days. Cells were also infected with empty vector (MigR1) and/or grown on control FC protein. **D.** PJ34 with CRISPR-mediated NOTCH1 KO were further engineered to express a DOX inducible cl-NOTCH cDNA (PJ34-iICN1) whose expression was titrated after 72 h treatment with different doses of DOX, to achieve protein levels equivalent to parental PJ34 stimulated with JAG1 for 16 hr. **E.** ICN1 induction with 1000 ng/ml DOX caused cells to massively shrink in size and form loosely attached tumor spheroids. **F.** Venn diagram illustrates overlap of genes in PJ34 significantly upregulated (FDR <0.1, |Fold change| ≥ 1.25-fold) by growth on JAG1 or after 24 h iICN1 induction, and genes specifically bound in the promoter or gene body regions by iICN1 after Chip-seq experiments. **G.** Venn diagram illustrating overlap of genes in PJ34 significantly downregulated (FDR <0.1, |Fold change| ≥ 1.25-fold) by growth on JAG1 or after 24 h iICN1 induction, and genes specifically bound in the promoter or gene body regions by iICN1 after Chip-seq experiments.

### Validation of Genes and Proteins Regulated by NOTCH1 Activation

Because of strong prior links between AXL or α-CATULIN and tumor aggressiveness and proliferation in HNSCC, we validated their decreased expression at the protein level following NOTCH1 activation in both *NOTCH1* WT and mutant tumors, using multiple approaches. First, we exogenously expressed activated cl-NOTCH1 using a retroviral construct encoding intracellular NOTCH1 (ICN1) widely used by others for functional studies (30). The ICN1 fragment encoded by this vector is missing several N-terminal amino acids at the cleavage site and is not recognized by antibodies specific for activated NOTCH1, but the integrity of the construct was confirmed through DNA sequencing and detection of its protein product after infecting *NOTCH1*-null mutant UMSCC22A with an antibody that binds the C-terminus of NOTCH1 (Supplementary Fig. S6A). Infection with ICN1 but not empty vector MigR1 induced morphological changes in UMSCC22A and *NOTCH1* WT 183 and PJ34 cells identical to that observed earlier for growth of these cell lines on JAG1 (Supplemental Fig. S6B, S6C, S6D). ICN1 expression caused substantial reduction in AXL and a-CATULIN proteins in *NOTCH1* WT PJ34 and 183 (Fig. 3B and Supplementary Fig. S6E) and *NOTCH1* mutant HN31 and UMSCC22A (Fig. 3B, and Supplementary Fig. S6A). Likewise, growth of PJ34, or *NOTCH1* mutant cells with restored NFL1 receptors (HN31, UMSCC22A, and UM47) on JAG1 ligand also suppressed AXL and a-CATULIN protein levels (Fig. 3C). Decreased LAMC2 and ITGA3 protein was also confirmed following ICN1 expression in PJ34 and HN31. Consistent with negative regulation by NOTCH1, increased AXL and α-CATULIN protein were found after pharmacological inhibition of NOTCH1 signaling with DBZ in the *NOTCH1* WT cell lines with normally high NOTCH1 activation (Supplementary Fig. S7).

### NOTCH1 Regulates Many Genes Indirectly

To better understand the mechanisms, timing and physiological relevance of NOTCH1 activation, we constructed a doxycycline (DOX) inducible ICN1 (iICN1) retroviral construct encoding intracellular NOTCH1 amino acids beginning from the known cleavage site that allowed more precise control over timing and levels of NOTCH1 activation, and could be detected by antibodies specific for cl-NOTCH1. We combined this tool with NOTCH1 Chip-seq experiments to identify genes directly regulated. *NOTCH1/NOTCH2* double knockout PJ34 cells (e.g., Supplementary Fig. S3) were engineered to express the Tet3 regulator (PJ34Tet3 cells) along with iICN1 (PJ34-iICN1) so that NOTCH1 was activated in the presence of DOX. Dose response experiments determined that 500-1000 ng/ml DOX induced levels of cl-NOTCH1 protein equivalent to those found after incubating parental PJ34 (PJ34-P) on JAG1 (Fig. 3D). Within one week after iICN1 induction with DOX, PJ34-iICN underwent the same morphological transformation observed earlier when parental PJ34 were grown on JAG1, characterized by massive cell shrinkage and formation of loosely attached tumor spheroids (Fig. 3E). RNA-seq performed from samples isolated 20 hours after peak iICN1 expression identified 1223 genes downregulated and 666 genes upregulated by 1.25-fold or greater (FDR <0.1) specifically in PJ34-iICN1 treated with DOX but not in control PJ34Tet3 cells treated with DOX (Table S7). NOTCH1 Chip-seq experiments identified 357 unique genes in PJ34-iICN1 bound by NOTCH1 at one or more loci in their gene promoters or gene bodies after DOX induction (Table S8). Venn diagrams illustrating the overlap of genes regulated by iICN1, JAG, and bound by ICN1 appear in Fig. 3F and 3G, with intersecting genes listed in Supplementary Tables S9 and S10. *AXL, α-CATULIN, ITGA3, ITGA5, LAMC1, and LAMC2* were all significantly downregulated by both iICN1 and JAG1 in PJ34, but only *LAMC2* was bound by ICN1 (e.g., within the gene body) in Chip-Seq experiments, suggesting the majority of observed changes linked to early differentiation were an indirect but early effect of NOTCH1 activation.

Genes from the HES (Hairy and Enhancer of Split) and Hey (Hairy/enhancer-of split with YRPW motif) family of transcriptional repressors are key canonical downstream targets of NOTCH1 signaling cascades that were also found to be elevated after ICN1 induction and identified through NOTCH1 Chip-seq. Specifically, HES2, HES4, HEY2, and HEYL were all bound by NOTCH1 in their promoter/gene body regions and significantly upregulated by 20h iICN1 induction (Supplementary Table S7 and S8) but showed lower fold changes with prolonged growth on JAG1 incubation (Supplementary Table S3 and S7). This likely reflects the cyclical nature of HES/HEY transcription following NOTCH1 activation. HES5, on the other hand, was bound by ICN1, and was strongly elevated after ICN1 induction or prolonged JAG1 exposure (Supplementary Table S3, S7 and S8). In contrast, the early differentiation markers *KRT13, KRT4*, and tumor suppressors *EPHA4, PDCD4,* and *TXNIP* ―all upregulated by prolonged JAG1 exposure― were not identified as direct NOTCH1 targets by Chip-seq nor were they strongly induced within 20 h of ICN1 expression (Supplementary Table S10), suggesting these are later events indirectly triggered through NOTCH1 signaling. Collectively the data support a model through which NOTCH1 activation triggers early but indirect suppression of cell adhesion receptors involved in transitioning away from basement membrane attachment with subsequent upregulation of differentiation markers.

### NOTCH1 Signaling Drives Anchorage Independent Growth but Fails to Increase Frequency of Cancer Stem Cells or Promote *In Vivo* Tumor Growth

*In vitro* growth of tumor spheroids in three-dimensional culture systems employing low-serum media is frequently used to propagate and quantify cancer stem cells (CSCs). The loosely attached tumor spheroids that formed in both *NOTCH1* mutant and *NOTCH1* WT cell lines after growth on JAG1 or NOTCH1 signaling stained positive for the senescence marker β-gal when cultivated in media with regular serum concentrations. However, serum is known to cause differentiation of CSCs. Therefore, we engineered some additional cell lines to robustly examine if NOTCH1 signaling would increase anchorage independent growth, survival, or expression of CSC markers in the presence of diminished serum, using our DOX iICN1 vector system. *NOTCH1*-null/mutant UMSCC22A cells were engineered to express the Tet3 regulator and iICN1. For FaDu, we used CRISPR gene KO to first delete endogenous *NOTCH1*, since baseline cl-NOTCH1 levels are normally high, before introducing the Tet3 regulator and iICN1. Titration experiments indicated that physiological levels of cl-NOTCH1 and the characteristic morphology changes and spheroid formation were achieved at doses of 250-500 ng/ml in FaDu-iICN1 (Fig. S8A and S8B). Physiological levels of activated NOTCH1 were achieved at 200-300 ng/ml DOX in UMSCC22A-iICN1, although morphology changes happened at an even lower dose (Supplementary Fig. S8C and S8D). At these doses of DOX, profound inhibition of growth in two-dimensional cultures accompanied expression of cl-NOTCH1 in both cell lines as well as PJ34-iICN1 (Supplementary Fig. S9A and S9B). In the absence of iICN1 infection, DOX failed to induce morphology changes or growth inhibition in any of the control Tet3G cells (Supplementary Fig. S9C). DOX-induced iICN1 expression significantly increased the number of tumor orospheres formed from FaDu and UMSCC22A (Supplementary Fig. S10A and S10B) in suspension cultures maintained with low serum, compared to control Tet3G cells or cultures lacking DOX.

Next, we examined whether increased tumor spheroid survival could be linked to increased resistance to anoikis (i.e., cell death associated with detachment). Tet3G control or iCN1cells were pretreated for 48 h with or without DOX and before trypsinization and incubation as single cell suspensions for an additional 30 hours in non-stick polypropylene tubes with constant rotation inside a 37 ^0^C tissue culture incubator. Induction of iICN1 was significantly protective in both FaDu and UMSCC22A compared to cells without DOX or control Tet3G cells in the presence or absence of DOX (Supplementary Fig. S10C). Because anoikis resistance and spheroid growth are both characteristics of CSCs, we examined whether NOTCH1 activation would also increase CSC markers previously associated with HNSCC, including Aldefluor activity, CD133 expression, and SOX2 protein levels. A 48-h induction of ICN1 failed to increase the percentage of CD44+bright/Aldefluor positive cells in both UMSCC22A and FaDu cells (Supplementary Fig. S11). Likewise, ICN1 induction failed to increase the percentage of cells expressing the CD133 surface marker or its mean fluorescence (Supplementary Fig. S12A and S12B). However, SOX2 gene expression was increased an average of 1.4-fold when PJ34 and 183 cells were grown on JAG and 1.3-fold in PJ34 after ICN1 induction (Supplementary Table S4 and S7), although SOX2 was not among genes bound by NOTCH1 in Chip-seq (Supplementary Table S8)―ruling out direct regulation. Likewise, levels of SOX2 protein were elevated after ICN1 induction in PJ34, FaDu, and UMSCC22A with increases ranging from 2- to 4-fold (Supplementary Fig. S12C).

Although stem cell markers and tumor spheroid formation can be surrogates for CSCs, the gold standard remains measuring the *in vivo* tumor initiating frequency in mice through limiting dilution assays. We reasoned that if NOTCH1 signaling were driving CSC behavior, it would most likely happen in *NOTCH1* WT tumors that endogenously express activated NOTCH1, like FaDu. Furthermore, for tumors to grow *in vivo*, stem cells must be allowed to re-enter a proliferative state resembling progenitors by turning NOTCH1 signaling off again. To avoid artifacts from non-physiological levels of NOTCH1 activation we conducted pilot studies to determine the *in vivo* DOX dose that would be equivalent to an *in vitro* dose of 400 ng/ml, which induced physiological levels of NOTCH1 signaling in FaDu-iICN1 (Supplementary Fig. S8A). Using a reporter cell line engineered to express luciferase from the same inducible promoter as iICN1 a dose of 1 mg DOX by oral gavage in mice led to equivalent fold induction of luciferase as 400 ng/ml DOX *in vitro* (Supplementary Fig. S13).

FaDu-iICN1 cells were then treated with 400 ng/ml DOX *in vitro* for 72h to activate NOTCH1 and potentially enrich for CSCs before inoculating increasing amounts of cells (100-100,000 cells range) subcutaneously into mouse flanks. This was followed by daily oral gavage with 1 mg DOX for an additional week to maintain NOTCH1 activation, followed by discontinuation of DOX for the remaining period to allow tumor cells to transition back to a proliferative state. As a control, matching numbers of untreated FaDu-iICN1 cells (e.g., no DOX) were inoculated into mice that never received DOX. Tumor cells treated with DOX grew much slower (Fig. 4A-D) and formed tumors later than tumor cells never treated with DOX at every inoculum dose (Fig. 4E-H). Eventually, tumors formed in 100% of animals for all groups except for the lowest inoculum of 100 cells, where only 40% of mice from the DOX group ever formed tumors by 100 days compared to 100% of mice that grew tumors within 40 days when no DOX was given. Statistical analysis estimated a tumor initiating frequency of 1/189 for DOX treated tumors compared to 1/1 for FaDu with no NOTCH1 induction (Table S11, P =0.00142), indicating a drastic reduction in CSC frequency with NOTCH1 activation.

**Figure 4.**
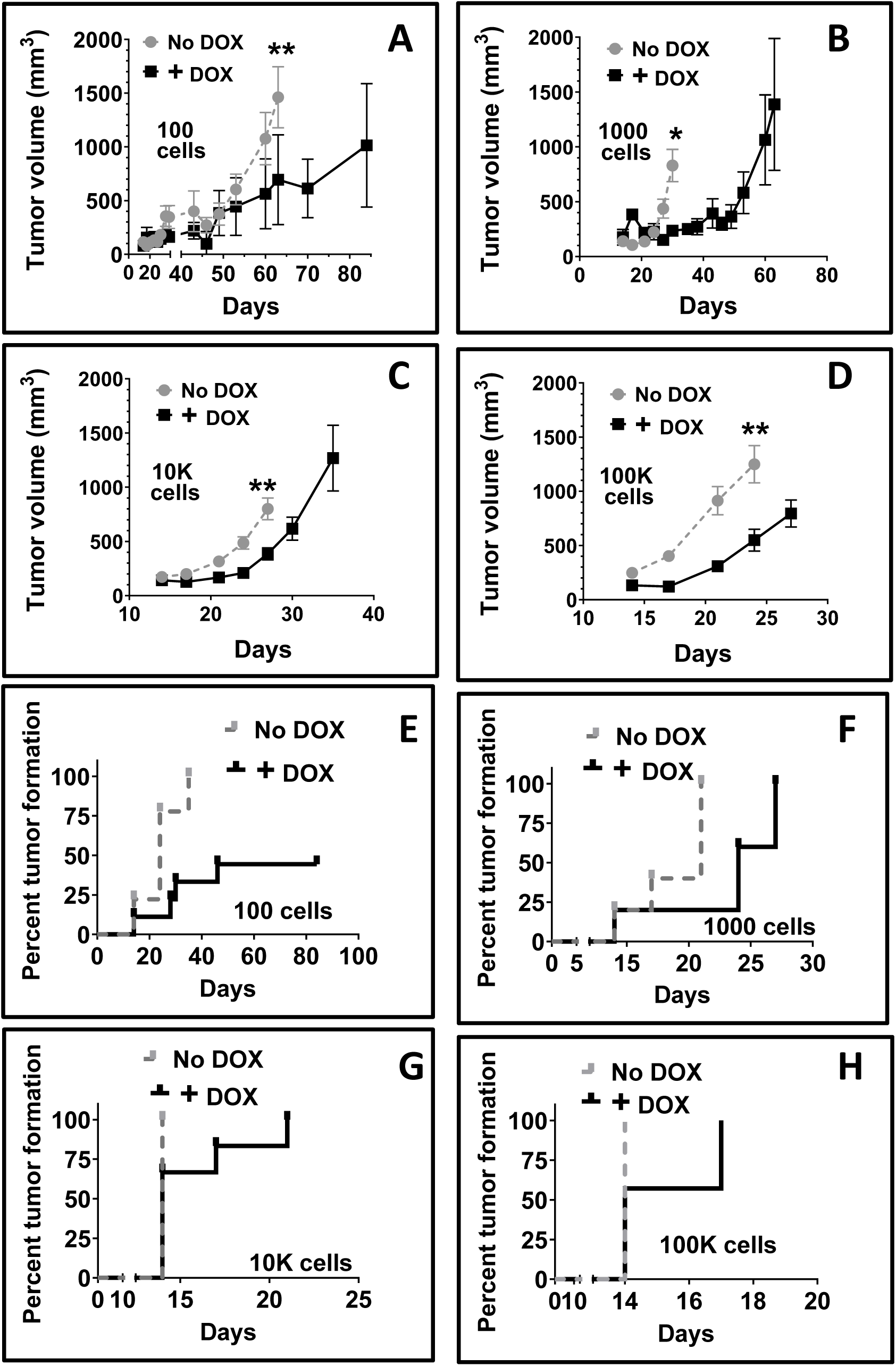
NOTCH1 activation reduces *in vivo* tumor growth and formation in NOTCH1 WT FaDU. After CRISPR-mediated NOTCH1 KO, FaDu were engineered to express iICN1 and pretreated with or without 300 ng/ml DOX for 72 hours *in vitro* before injecting 100 cells (A), 1000 cells (B), 10,000 cells (C), or 100,000 cells (D) into flanks of mice. The DOX treated group received additional *in vivo* DOX (1 mg) for one week by oral gavage after implantation and tumor growth was plotted verses post-inoculation time. Time to tumor formation in mice was plotted verses post-inoculation time for 100, 1000, 10,000, or 100,000 inoculated tumor cells (E-F) in the no Dox (grey) or DOX (black) treated groups.

When NOTCH1 signaling was restored in a *NOTCH1* mutant background using UMSCC22A-iICN, it also profoundly inhibited tumor growth. UMSCC22A-iICN1 were implanted into flanks of nude mice, which were randomized to receive placebo or 1 mg DOX by oral gavage daily for 3 weeks to persistently induce iICN1. NOTCH 1 activation profoundly suppressed *in vivo* tumor growth (Supplementary Fig. S14A). After discontinuation of DOX, tumors eventually grew in some mice. Histological staining revealed a roughly 50% reduction in the presence of mouse fibroblasts within the DOX treated tumor group (P<0.01, Supplementary Fig. S14B).

### Suppression of Oncogenic AXL and α-CATULIN Expression Contributes to NOTCH1-mediated Growth Inhibition

NOTCH1 activation increased expression of multiple tumor suppressor genes and simultaneously reduced expression of AXL and α-CATULIN, two oncogenic proteins linked to tumor growth and aggressiveness in HNSCC. We functionally examined the impact of reduced *AXL* and *α-CATULIN* expression on tumor growth *in vitro* and *in vivo*, with bicistronic shRNA IRES EGFP constructs targeting these two genes. Phenotypes were measured shortly after purifying infected (e.g., EGFP+) cells. Specific knockdown of either AXL or α-CATULIN protein in both *NOTCH1*-mutant HN31 and *NOTCH1*-WT PJ34 was confirmed by Western blotting (Supplementary Fig. S15A and S15B) and found to significantly reduce colony formation in clonogenic assays (P<0.006, Supplementary Fig. S15C and S15D). Importantly, shRNA knockdown of either *AXL* or *α-CATULIN* severely diminished tumor growth of HN31 in an orthoptic tongue tumor model (Supplementary Fig. S15E) and in a subcutaneous flank model using alternate shRNA sequences (Supplementary Fig. S15F), demonstrating the phenotype was robust.

We used PJ34-iICN1 and UMSCC22A-iICN1 to functionally examine whether preventing NOTCH1-induced increases in HES/HEY family members would prevent associated decreases in *AXL* and *α-CATULIN* gene expression. In both cell lines, prior knockdown of *HES1, HES2, HES4, HEY1,* or *HEY2* with siRNA shortly before ICN1 induction reduced their elevation stemming from NOTCH1 activation but did little to prevent associated reductions in *AXL* and *α-CATULIN* expression (determined by qPCR, Supplementary Fig. 16). Furthermore, combined simultaneous knockdown of some of the more strongly induced NOTCH1 targets, *HES5/HEY1/HEY2*, also failed to prevent NOTCH1-induced decreases in *AXL* and *α-CATULIN* expression in both PJ34 and UMSCC22A (Not shown).

### A Gene Expression Signature identifies Primary HNSCC Tumors with Intact NOTCH1 Signaling and an Altered Tumor Microenvironment

A robust *in vivo* gene expression signature of NOTCH1 activation was developed based on the top 120 differentially regulated genes identified *in vitro* after JAG1 stimulation (Supplementary Fig. S5) by examining their cross-correlation in primary tumors from The Cancer Genome Atlas (TCGA) oral squamous cell carcinoma (OSCC) cohort of 312 patients. After removing one gene for low expression, two-way hierarchical clustering of cross-correlation coefficients identified two primary gene clusters or modules (Supplementary Fig. S17A) from the TCGA data. Genes from the two clusters were annotated according to their *in vitro* regulation and it was clear that each cluster had a dominant but opposite pattern of *in vitro* gene regulation by JAG1, which identified groups of genes that were up or down regulated. After removal of several inconsistently regulated genes, 95 genes remained in the final signature (Supplementary Fig. S17B and Table S12). We applied the 95 gene signature to perform consensus hierarchical clustering on the TCGA OCSCC and TCGA laryngeal\hypopharyngeal SCC (LHSCC) cohorts individually. Consensus clustering metrics identified that the choice of two sample clusters was optimal for both TCGA cohorts (Supplementary Fig. S18). After two-way clustering (Fig. 5A and 5B), it was predicted that NOTCH1 signaling is off in sample cluster 1 but on in sample cluster 2 for both OCSCC and LHSCC cohorts, based on the direction of gene regulation from the 95-gene signature observed (Tables S13-14). The predicted NOTCH1 pathway states were compared to the *NOTCH1* mutational status of samples in both cohorts to validate the NOTCH1 signature. In both cases, sample cluster 2 was depleted for *NOTCH1* mutations but sample cluster 1 was enriched (P = 0.015 and P = 0.013, Fig. 5A and B), consistent with inactivating *NOTCH1* mutations preventing NOTCH1 pathway signaling. Consequently, the gene signature likely distinguished tumors based on their NOTCH1 pathway status, and the data suggests that NOTCH1 signaling is active in a subset of OCSCC and LHSCC tumors (e.g., sample cluster 2).

**Figure 5.**
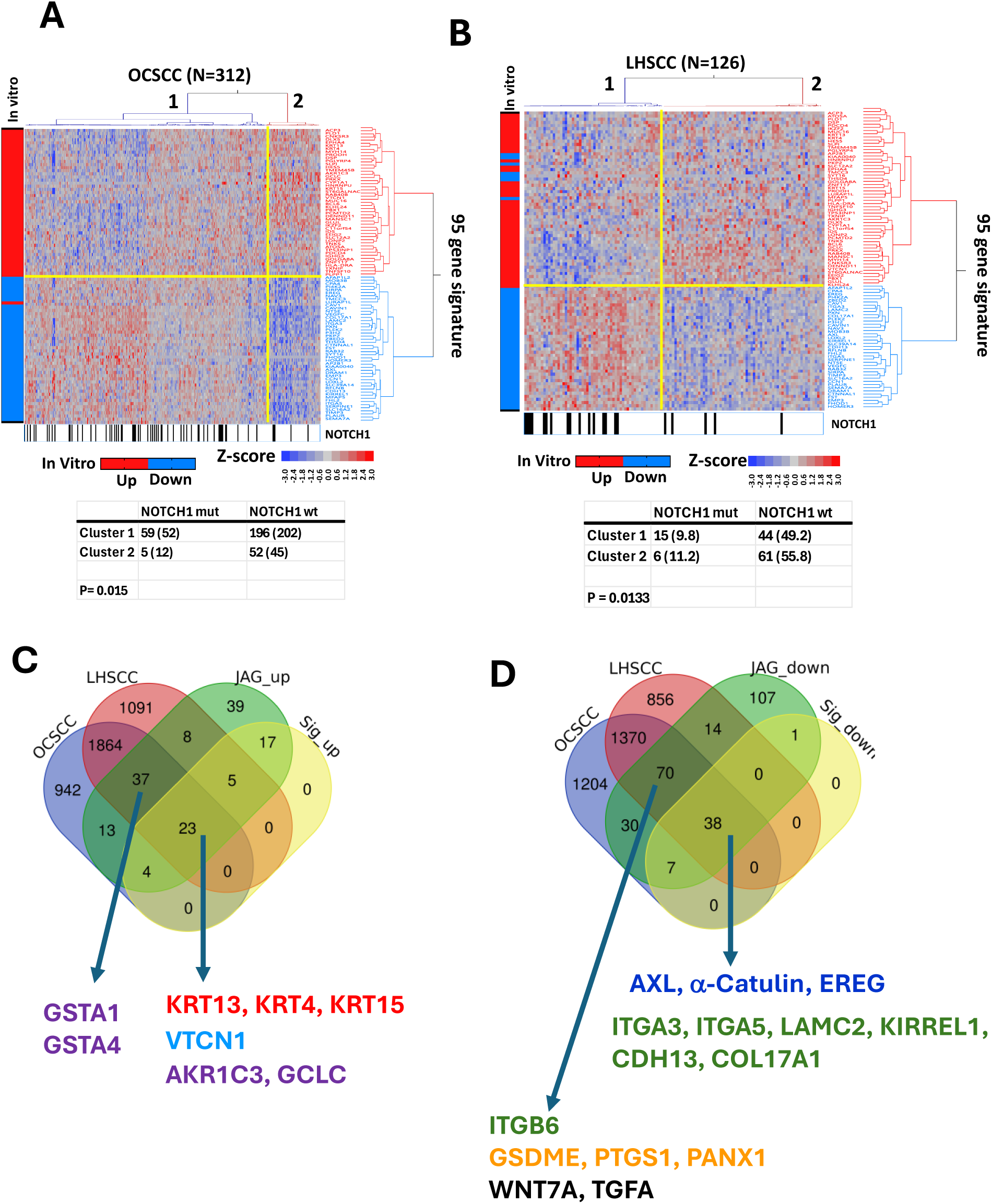
Comparison of genes regulated by NOTCH1 *in vitro* and genes differentially expressed in primary HNSCC tumors with a NOTCH1 activation signature. **A**. Consensus hierarchical clustering of TCGA OCSCC specimens based on a 95-gene NOTCH1 activation signature identified a cluster of patients (Cluster 2, N = 57) with an expression pattern indicative of active NOTCH1 signaling and another cluster (Cluster 1, N = 255) predicted to have loss of NOTCH1 signaling. Genes are annotated with vertical boxes according to whether they were upregulated (red) or downregulated (blue) by JAG1 in vitro. Samples with a NOTCH1 mutation are annotated horizontally with a black box and association between NOTCH1 mutation status and cluster for patients with sequencing information was analyzed by Chi-square analysis. **B.** Parallel clustering and analysis of TCGA LHSCC samples using the same 95-gene NOTCH1 signature. **C.** Venn diagram illustrating overlap of genes upregulated (FDR<0.1, |FC|≥1.25) by NOTCH1 in vitro (JAG_up), the subset of upregulated genes part of the NOTCH1 signature (Sig_up), and genes upregulated (FDR<0.1, |FC|≥1.25) in Cluster 2 from OCSCC or LHSCC. **D.** Venn diagram illustrating overlap of genes downregulated (FDR<0.1, |FC|≥1.25) by NOTCH1 *in vitro* (JAG_down), the subset of downregulated genes part of the NOTCH1 signature (Sig_down), and genes downregulated (FDR<0.1, |FC|≥1.25) in Cluster 2 from OCSCC or LHSCC.

Next, we identified *all genes* differentially expressed by the subset of TCGA tumors predicted to have NOTCH1 activated in the OCSCC (Table S15) and LHSCC (Table S16) cohorts and compared them to genes found differentially regulated by JAG1 binding *in vitro*, including the subset of 95 genes defining the pathway signature. Venn diagrams depicting overlap of differentially expressed genes appear in Fig. 5 C and D. Among the commonly *upregulated* genes (Supplementary Table S17) were early differentiation markers ***KRT13***, ***KRT14**, and **KRT15***; the immune checkpoint **VTCN1**; and several enzymes involved in the antioxidant response, including ***AKR1C3***, which neutralizes lipid peroxides, as well as ***GCLC***, ***GSTA1***, and **GSTA4**, which are essential for glutathione synthesis and anti-oxidant response. Among commonly downregulated genes (Supplementary Table S18) were multiple integrins and cell adhesion molecules ***ITGA3, ITGA5, ITGB6, LAMC2, KIRREL1, CDH13,*** including ***COL17A1*** (collagen XVII) that mediates adhesion to basement membranes as a component of hemidesmosomes. The pro-oncogenic genes ***AXL*** and ***α-CATULIN*** were also commonly downregulated. So was ***WNT7A***, which contributes to HNSCC growth by stimulating β-catenin, as well as ***EREG*** and ***TGFA***, both EGFR ligands that promote HNSCC.

<u>S</u>ingle <u>s</u>ample <u>G</u>ene <u>S</u>et <u>E</u>nrichment <u>A</u>nalysis (ssGSEA) was used to determine associations between the tumor microenvironment and NOTCH1 signaling, using published lists (Table S19) specific for immune subtypes, endothelial cells, and a robust gene list we constructed for cancer associated fibroblasts (CAFs). Our CAF signature was derived by clustering cross-correlation coefficients for fibroblast associated genes across more than 9000 solid tumors from the TCGA. In the TCGA OCSCC cohort, tumors with a NOTCH1 activation signature had significantly reduced proportions of nearly every immune subset analyzed (Table S20), indicating they are immunologically “cold”. Two-way hierarchical clustering with the immune subset ssGSEA scores demonstrated significant depletion of tumors with active NOTCH1 signaling among OCSCC “hot” tumors (P<0.0001, Fig. 6A). In LHSCC samples with NOTCH1 activation there was a similar trend of broad decrease in leukocytes subpopulations present when NOTCH1 signaling was on, although differences only reached statistical significance for T helper type 1 and mast cells (Table S21). Because we previously showed that a cold immune microenvironment was associated with elevated NRF2 gene signatures specifically in OCSCC but not LHSCC (31), we revisited the connection between NOTCH1 signaling and the antioxidant response inferred from some of the top genes commonly regulated by NOTCH1 (Fig. 5C) or proteins correlating with cl-NOTCH1 (Table S2). HNSCC TCGA tumors with NOTCH1 activation had NRF2 pathway scores (Fig. 6B) that were on average profoundly elevated regardless of disease subsite (P<0.0001). In contrast, NOTCH1 activation was associated with a significant reduction in the proportion of CAFs present, imputed from CAF ssGSEA scores in both OCSCC and LHSCC (Fig. 6C), which was supported by gross differences in fibroblast content visible in H&E images downloaded from the TCGA project (Supplementary Fig. S19).

**Figure 6.**
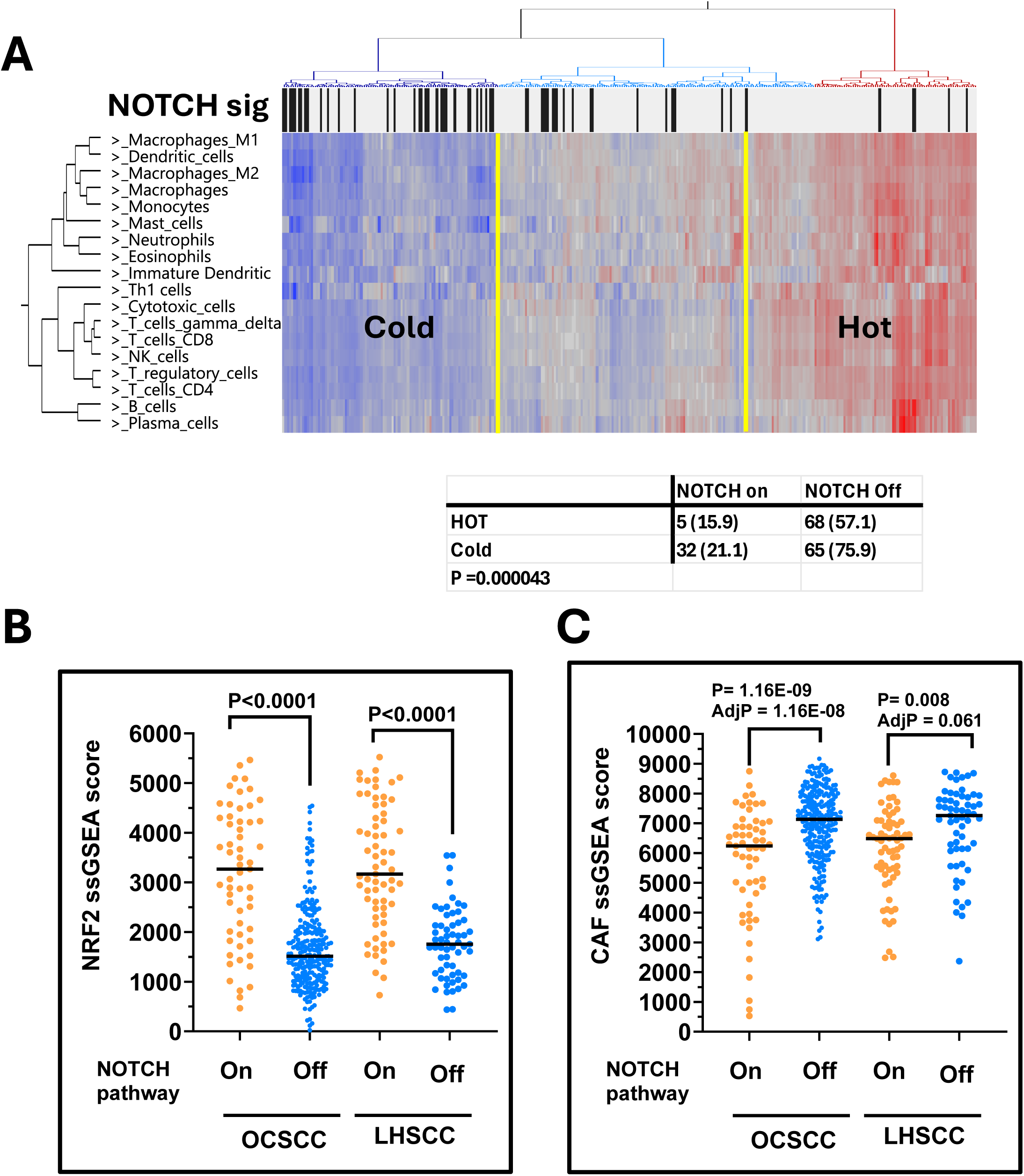
NOTCH1 activation in primary HNSCC is associated with changes to the tumor microenvironment. **A.** ssGSEA scores representing 18 different leukocyte subsets derived from TCGA OCSCC were used for hierarchical clustering to classify samples as immunologically cold (sample cluster 1) or hot (sample cluster 3) and the membership of samples from previous clustering based on the NOTCH1 gene signature is annotated with a black box for NOTCH1 signaling on or a grey box for NOTCH1 signaling off. **B.** NRF2 pathway scores are compared across NOTCH1 sample clusters from OCSCC and LHSCC TCGA tumors. **C.** CAF pathway scores, indictive of CAF infiltration, are compared across NOTCH1 sample clusters from OCSCC and LHSCC TCGA tumors.

### NOTCH1 Signaling Correlates with Survival and *PIK3CA* Genomic Alterations

When NOTCH1 signaling status was treated as a dichotomous variable for OCSCC TCGA samples based on clustering, there was no significant association (Fig. 7A) with overall survival (OS). Because signaling pathways are rarely binary, we used the NOTCH1 gene signature to generate ssGSEA scores for samples and treated NOTCH1 signaling as a continuous variable, which demonstrated clear separation between sample clusters found previously (Fig. 7B). Recursive portioning identified a NOTCH1 ssGSEA threshold of −1554 that stratified OCSCC patients into two groups which differed significantly by survival with poor survival corresponding to lower levels of NOTCH1 signaling (P = 0.0061, Fig. 7C). The threshold identified seemed biologically meaningful as it corresponded to roughly the average value for samples from the NOTCH1-off clusters in both OCSCC and LHSCC (Fig. 7B), and it also separated the LHSCC samples into two groups that differed in survival in the same manner (P = 0.0.0045, Fig. 7D). Likewise, patients with NOTCH1 ssGSEA scores below this same threshold had significantly worse progression free survival in both disease subsites (Fig. 7E and F). No associations were found between lymph node stage, tumor stage, or smoking history in either disease subsite, although there was a significant decline in NOTCH signaling levels among poorly differentiated tumors found in OCSCC tumors (not shown). We previously reported that HNSCC cell lines harboring *NOTCH1* loss of function mutations are exquisitely sensitive to PI3K inhibitors (27,28), which was supported by a small clinical trial we conducted (29). In those studies, we found direct links between NOTCH1 signaling itself and PI3K inhibitor sensitivity to be nuanced and context dependent, which led us to hypothesize that tumors evolving with *NOTCH1* mutations may have altered pathway dependencies, including PI3K. We analyzed the relationship between TCGA NOTCH1 sample clusters and found that the presence of PIK3CA genomic alterations (e.g., high level copy gains or mutations) was significantly enriched among OCSCC (P = 0.024) and LHSCC samples (P = 0.0004) belonging to the NOTCH-On clusters (Supplementary Fig. S20).

**Figure 7.**
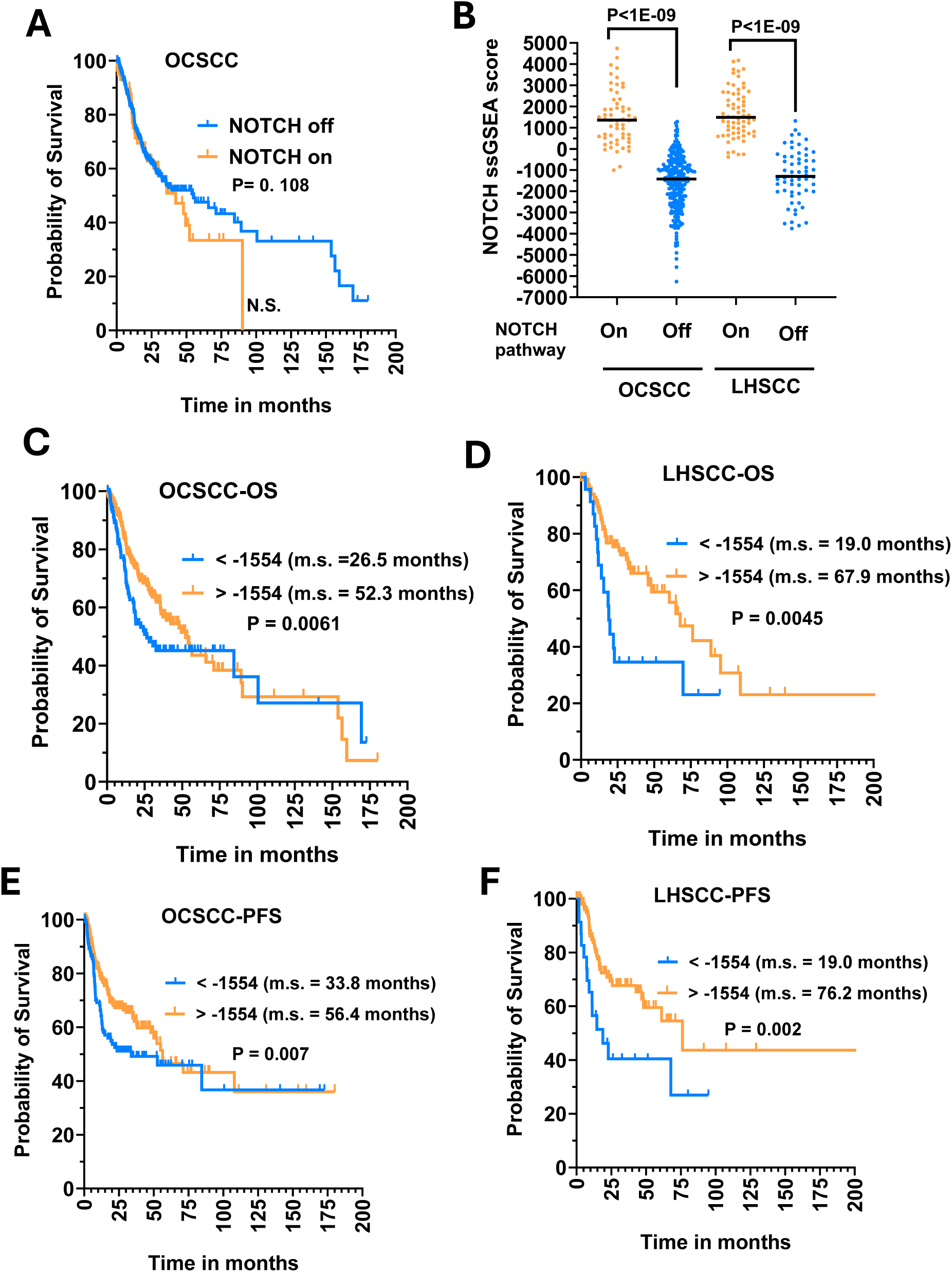
Higher levels of NOTCH1 activation correlate with better survival in OCSCC and LHSCC TCGA cohorts. **A.** No difference in OS among TCGA OCSCC patients when NOTCH1 activation is treated as a categorical variable based on clusters with the NOTCH gene signature. **B**. Validation that ssGSEA scores derived from the NOTCH1 gene signature provide a continuous value measurement that faithfully replicates sample clustering. **C.** OCSCC samples with higher ssGSEA scores above a threshold (−1554) determined by recursive partitioning have significantly improved OS. **D.** LHSCC samples with higher ssGSEA scores above the same threshold (−1554) have significantly improved OS, validating the threshold. **E.** OCSCC patients with higher NOTCH1 signaling have improved PFS. **F.** LHSCC patients with higher NOTCH1 signaling have improved PFS.

## DISCUSSION

More than a decade has passed since our group and others first identified inactivating *NOTCH1* mutations as a driver of HNSCC (1,3). Yet, the function and significance of NOTCH1 signaling in this cancer are still poorly understood. Adding to the complexity, various reports suggest that WT *NOTCH1* acts as an oncogenic driver in some HNSCC cell lines by inducing CSC-like properties. Additionally, strong NOTCH1 signaling has been observed in subsets of primary HNSCC tumors (7,26). In the present study, we clarify the biology of NOTCH1 signaling in HNSCC and unify seemingly conflicting schools of thought.

Regardless of *NOTCH1* mutational status or endogenous levels of signaling, the pathway activates a program of very early differentiation that involves downregulation of cell adhesion molecules, which normally tether cells to their basement membrane, accompanied by upregulation of keratin differentiation markers. These changes in adhesion accompanied by increased anoikis resistance, which is likely a vestige of squamous epithelial stratification, promotes growth of cells in a non-adherent spheroid state and likely explains reports that NOTCH1 signaling triggers HNSCC tumors to become more CSC like (19,25). However, using an inducible cl-NOTCH1 expression system in *NOTCH1* WT FaDu clearly showed a substantial decrease in tumor initiating cells *in vivo* using the gold standard limiting dilution assay. Furthermore, we found that a NOTCH1 inhibitor DBZ, more potent and specific than DAPT frequently used in prior NOTCH studies (16,23,32), had no effect on *in vitro* growth of six different HNSCC tumor lines specifically chosen for high basal cl-NOTCH1expression. Collectively, this refutes the idea that NOTCH1 actively drives cell proliferation in HNSCC. Interestingly, we find evidence of limited NOTCH1 signaling in HNSCC cell lines with high basal cl-NOTCH1 expression, given the strong anti-correlation with AXL protein―one of the genes robustly downregulated by NOTCH1 activation. That said, no obvious growth phenotypes distinguished cell lines with high NOTCH1 signaling but when the gene was knocked out in FaDu and cl-NOTCH1 re-expressed at physiological levels there was a dramatic morphologic transformation to tumor spheroid growth not otherwise observed for parental FaDu with equivalent NOTCH1 signaling. Expression of Fbxw7 protein, the known E3-ligase to degrade intracellular activated NOTCH1, strongly correlated with cl-NOTCH1 protein by RPPA in cell lines. Collectively, our data suggest that some HNSCC cell lines can tolerate NOTCH1 signaling *in vitro* and that growth in two-dimensional cultures may select for diminished downstream phenotypes.

By defining genes altered after physiological activation of NOTCH1 *in vitro*, we were able to construct an empirically derived gene expression signature of NOTCH1 activation that was validated in primary HNSCC tumors from two different disease sites, allowing stratification of patients’ samples based on relative levels of NOTCH1 signaling. Wholesale downregulation of cell adhesion receptors, particularly integrins along with genes encoding extracellular matrix proteins such as laminins and COL17A1 were found to accompany NOTCH1 activation in both preclinical models and primary tumors, consistent with very early differentiation of tumors that mirrors transition from the basal to suprabasal layer in normal mucosa. Possibly, the shift in cell adhesion and extracellular matrix proteins could impact the tumor microenvironment and contribute to the diminished presence of CAFs. We found evidence of this in our preclinical model and in both disease subsites when NOTCH1 signaling was elevated. A surprising novel finding was the association between NOTCH1 activation and increased NRF2 pathway signaling, which has implications for chemoradioresistance, and is consistent with reports that NOTCH1 signaling increases chemotherapy resistance (25). We previously published that elevated NRF2 activity contributes to cisplatin resistance using preclinical models (33,34) and that elevated NRF2 activity is associated with an immunologically cold tumor immune microenvironment in multiple tobacco associated tumors (31), including OCSCC but not LHSCC. Consistent with this, OCSCC tumors with elevated NOTCH1 signaling had increased NRF2 scores and broadly diminished infiltration of leukocyte subtypes.

Many genes altered by NOTCH1 were modulated within an early time frame but were not directly regulated by binding of intracellular NOTCH1 to their promoters or enhancers. Of these, *AXL* (35,36) and *α-CATULIN* (37,38) reportedly contribute to HNSCC tumor growth in preclinical models and are associated with clinical aggressiveness. We confirmed their pro-oncogenic function *in vitro* and in *vivo* through knockdown experiments and they contributed to some of the growth inhibition triggered by NOTCH 1 activation. However, the multiplicity of genes commonly regulated by NOTCH1 both *in vitro* and *in vivo* suggests cooperative gene expression programing with possibly redundant function. SOX2 was the one gene tied to CSC that we and others found upregulated by NOTCH1 activation (24,39). However, IHC has shown that SOX2 (40) along with cl-NOTCH1 (7) is frequently present in the normal mucosal suprabasal layer and elevated SOX2 correlates with better HNSCC prognosis (40), consistent with our proposed model of very early differentiation.

If NOTCH1 signaling turns on a program of very early differentiation and is not associated with CSC maintenance, then we might expect tumors with higher NOTCH1 signaling to have a better prognosis. Consistent with what others reported for cleaved NOTCH1 staining (26), we found that subsets of tumors with higher NOTCH1 signaling scores had improved OS and PFS across disease subsites. Our preclinical models demonstrated that WT *NOTCH1* tumors retain plasticity and can undergo very early differentiation in response to NOTCH1 signaling. This is consistent with prior work proposing that *NOTCH1* LOF mutations drive carcinogenesis by preventing early stem cell differentiation and promoting accumulation of secondary mutations in an expanding pool of stem cells (41). Possibly, precancerous lesions arising with *NOTCH1* mutations avoid early differentiation and retain some pathway dependencies of stem cells that may include the PI3K pathway, which has been linked to survival and maintenance of CSCs (42). This could explain the sensitivity to PI3K inhibitors associated with *NOTCH1* mutations we reported in the absence of *PIK3CA* mutations (27). If early differentiation driven by NOTCH1 activation in WT tumors were accompanied by a shift away from the PI3K pathway, then perhaps these tumors rely more on genomic alterations in the *PIK3CA* gene to re-establish signaling. This is supported by our findings that mutations and high-level amplifications of the *PIK3CA* gene are more frequent in tumors with higher NOTCH1 signaling.

How then do we reconcile our conclusions with multiple reports of NOTCH1 behaving like an oncogene in HNSCC? Many factors likely contribute, including overinterpretation of tumor spheroid properties in this context, frequent use of antibodies that detect inactive membranous/cytoplasmic NOTCH1 rather than active cleaved NOTCH1 for IHC studies, nonspecific pharmacological inhibitors to block NOTCH1 signaling, widely available constructs to overexpress activated NOTCH1 that cannot be validated with cleavage specific antibodies, and/or ambiguous use of antibodies recognizing C-terminal NOTCH1 peptides in lieu of those specific for activated NOTCH1. In summary, we find molecular evidence of NOTCH1 signaling in subsets of HNSCC tumors that has broad gene expression consequences impacting tumor biology, the tumor microenvironment and clinical behavior. However, most of the evidence unequivocally supports a tumor suppressor function for *NOTCH1,* regardless of mutational status or baseline NOTCH1 signaling, which triggers early differentiation accompanied by decreased cell attachment.

## METHODS

### Cell lines, plasmids, and reagents

The established HNSCC cell lines used in experiments and listed in Supplementary Table S1 were passaged in growth media containing 10% FBS plus additives, validated by STR profiling, and profiled for somatic mutations as previously described (43). Full-length WT human *NOTCH1* receptor (NFL1) cDNA was obtained from OriGene (Rockville, MD), the ICN1 retroviral construct encoding activated human *NOTCH1* from Dr. Patrick Zweidler-McKay, and the cDNA encoding human cleaved *NOTCH1* detectable by commercial cl-NOTCH1 antibodies was subcloned through reverse transcriptase PCR using RNA derived from *NOTCH1* WT FaDu cells. Details regarding these constructs, shRNAs targeting *AXL* and *a-CATULIN* and CRISPR vectors, and siRNA reagents targeting *HES/HEY* family members are provided in supplementary methods. Catalogue numbers for antibodies obtained from Cell Signaling Technologies (Danvers, MA) and Santa Cruz (Santa Cruz, CA) are provided in supplementary methods as well.

### NOTCH Activation, Clonogenic Assays, Western Blots, Reverse Phase Protein Arrays

For NOTCH activation experiments, tissue culture wells were pre-coated overnight at room temperature (RT) with Protein G (Prospec, East Brunswick, NJ) at 50 ug/ml, washed twice in phosphate buffered saline (PBS), blocked in 1% BSA/PBS for 2 hours at RT, washed three times with PBS, coated with either human recombinant chimeric Jag1 fused to an FC fragment (R&D Systems, Minneapolis, MN) or control purified IgG FC protein (Jackson ImmunoResearch, West Grove, PA) at 2 ug/ml in 0.1% BSA/PBS for 3 hours RT, stored overnight at 4 ^0^C, and washed three times immediately before use. Proteins lysates were harvested, resolved by SDS-PAGE electrophoresis, electro-transferred to PVDF membranes, and probed with specific antibodies using standard methods as previously described. Antibodies to activated cleaved NOTCH1 (cl-NOTCH1), total NOTCH1, total NOTCH2, AXL, FLAG tag, were from Cell Signaling; whereas antibodies to α-CATULIN LAMC2, ITGA3, ITGA5 were from Santa Cruz and anti-Hes5 was from Abcam. For clonogenic assays, 1000 cells were seeded into replicate 6-well plates either uncoated or pretreated with Jag1 or control FC, grown for 7-10 days, fixed and stained with 0.5% crystal violet, and the number of colonies with greater than 60 cells were counted with Image J software (3). RPPAs were used to quantitate levels of 157 different protein/phosphoproteins using lysates prepared from a panel of HNSCC cell lines and validated antibodies according to methods we previously published.

### Differentially Expressed Gene (DEG) Analysis and RNA-seq

After growing cells on plates coated with either Jag1 or FC control protein (in biologic triplicate) for 5 days, extracted RNA was processed by the MD Anderson Sequencing and Microarray Core Facility, where hybridization of replicates was individually quantitated following hybridization to Affymetrix HuGene 2.0 ST arrays. RNA expression after induction of iICN1 in PJ34-iICN1 cells was determined by RNA-seq using replicate samples incubated in the absence or presence of 1000 ng/ml DOX for 36 h. Detailed bioinformatics analyses, including identification of DEGs, consensus hierarchical clustering, and comparison to the HNSCC TCGA RNA-seq data from 423 HNSCC patients are described in supplementary methods.

### Quantitative Real Time PCR (qPCR)

RNA (2-5 μg) was converted to cDNA using a SuperScript First Strand synthesis Kit (Life Technologies) and 100 ng of cDNA was added to quadruplicate reactions containing FAM-MGB TaqMan PCR primers (supplementary methods) and amplified in a c1000 Bio-Rad Thermal Cycler. Expression of targets was normalized to GAPDH by the ΔCT method using CFX Manager 3.1 software (BioRad).

### Staining for β-galactosidase

Cells were fixed and stained with a β-Galactosidase Kit (Cell Signaling), according to supplied instructions and 5 random 10x objective fields were observed to count the number of positive and negative staining cells. For quantitating spheres, 25 random fields were counted.

### Mouse Tumor models and Cancer Stem Cell Frequencies

Cells infected with equivalent titers of shRNA lentivirus (*AXL*, *α-CATULIN*, or empty vector) were sorted for GFP positivity by flow cytometry, allowed to recover for 2 days and then 50,000 cells (in 30 μl PBS) were injected into the anterior tongues of anesthetized male nude mice (10 per group) using a 30-gauge needle. For flank models utilizing tumors infected with shRNA, 3-4 million cells were injected subcutaneously in 200 μl PBS. For *in vivo* experiments utilizing FaDU-iICN1 or UMSCC22a-iICN1, 3-4 million cells were injected subcutaneously into flanks of nude mice. For some groups, mice were administered 1 mg doxycycline (DOX) dissolved in 200 μl water by oral gavage 5 days per week, for 1 to 3 weeks as described in the text. Cancer stem cell frequencies in untreated and DOX treated FaDu-iICN1 populations were estimated and compared from results of limiting dilution experiments using the Extreme Limiting Dilution Analysis Software tool (44) at https://bioinf.wehi.edu.au/software/elda/.

Mouse manipulations were performed in accordance with a protocol approved by an institutional animal care and use committee.

## Data availability

The data generated in this study are available in the article and its supplementary files.

**Supplementary Figure 1.**
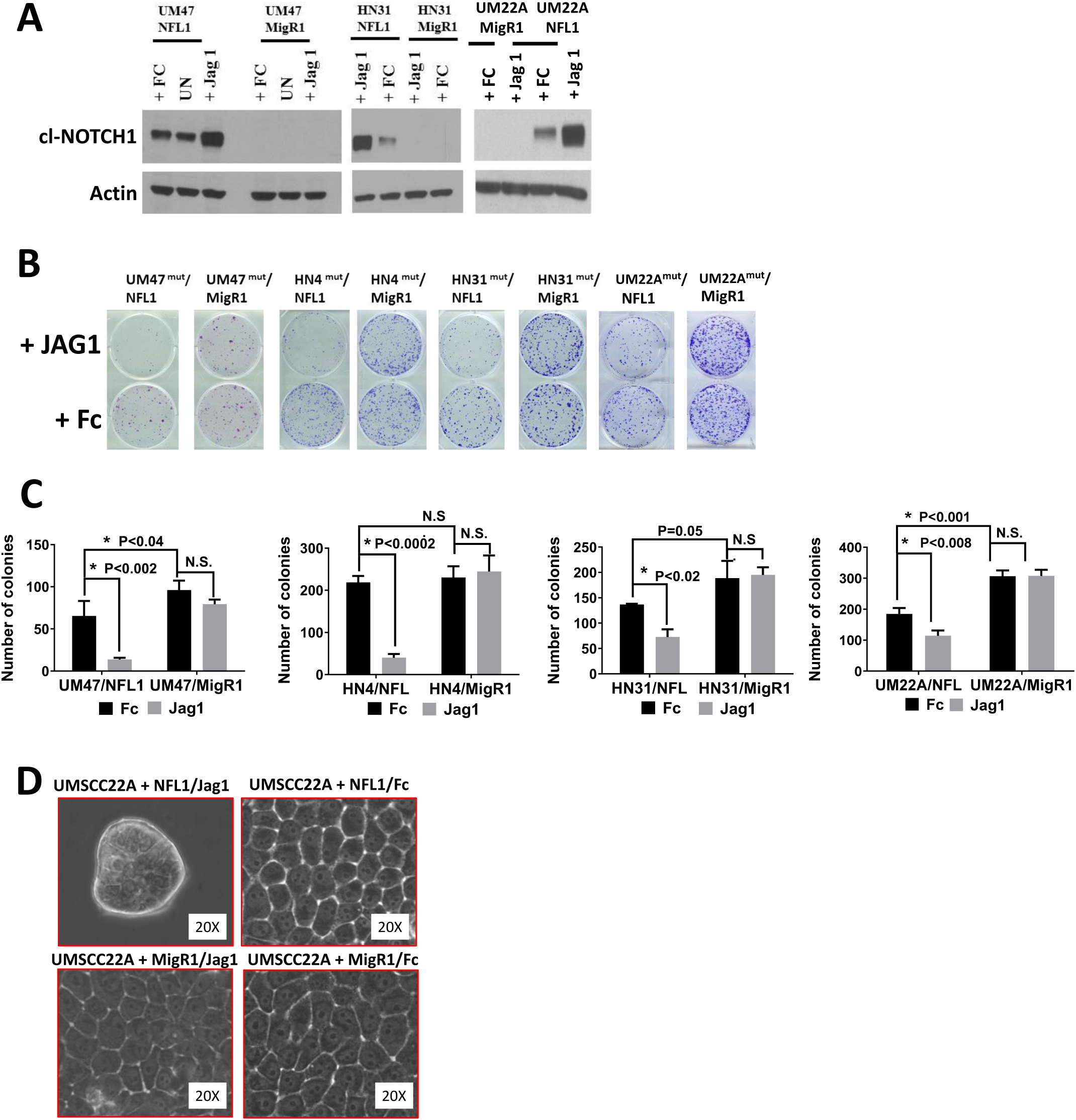
Restoration of NOTCH1 signaling inhibits *in vitro* growth and alters morphology in NOTCH1 mutant HNSCC tumors. **A**. Ectopic expression of WT NFL1 leads to activation and cleavage of NOTCH1 (Cl-NOTCH1) protein in 3 different NOTCH1-mutant HNSCC cell lines that increases upon 16 h stimulation with immobilized JAG1 ligand, compared to control FC protein. **B**. Continued growth on immobilized JAG1 for 8-12 days inhibits colony formation in NOTCH1-mutant cell lines with restored NFL1. **C**. Quantitation of NFL1-mediated growth inhibition of colonies. **D.** Restoration of NFL1 signaling in UMSCC-22A led to dramatic reduction in cell size and formation of loosely attached tumor spheroids after 5 days only in the presence of JAG1. UM47 = UMSCC47, UM22A = UMSCC22A

**Supplementary Figure 2.**
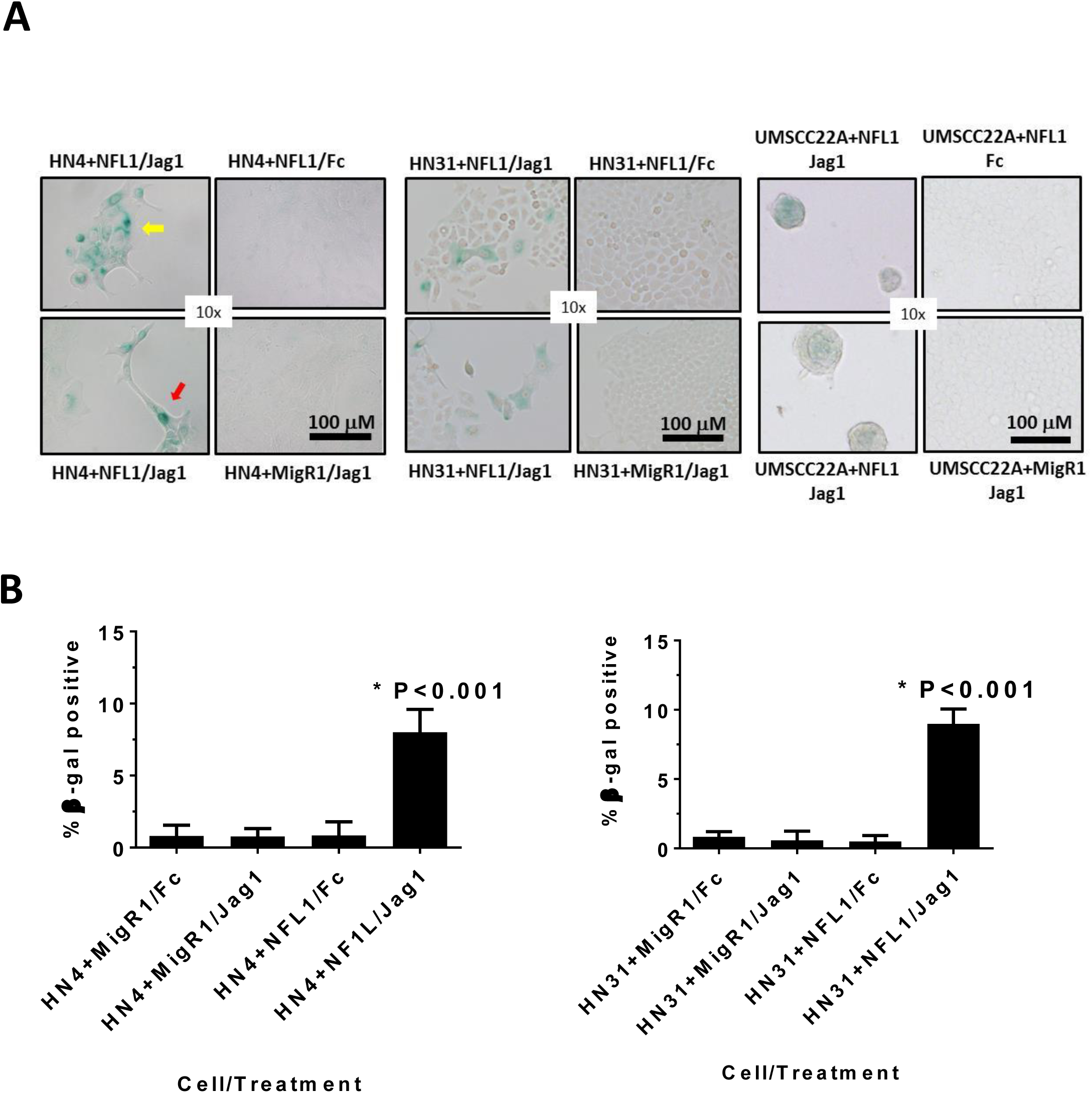
Restoration of NOTCH1 signaling induces senescence. **A.** Photomicrographs of *NOTCH1*-mutant HNSCC cell lines infected with NFL1 or empty MigR1 grown on either JAG1 or FC control protein and stained for β-gal after 5 to 7 days culture. **B.** Quantitation shows significant elevation of β-gal staining in the presence of NFL1 expression and growth on JAG1 ligand.

**Supplementary Figure 3.**
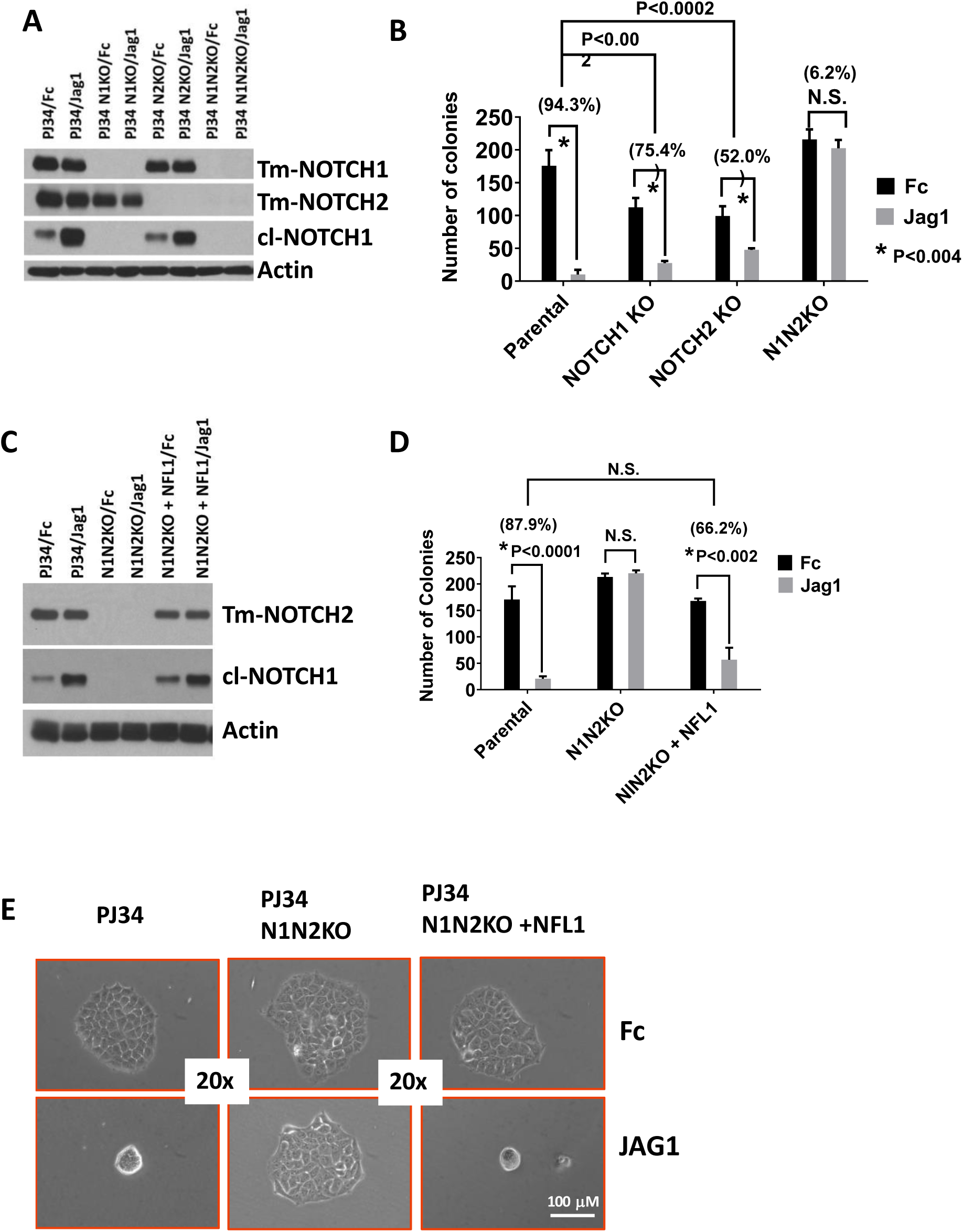
Both NOTCH1 and NOTCH2 signaling contribute to JAG1-induced growth inhibition but NOTCH1 signaling is sufficient. **A.** CRISPR KO of either *NOTCH1* (NIKO), NOTCH2 (N2KO) or double KO (N1N2 KO) in PJ34 were validated by western blotting for total NOTCH1 and NOTCH2. Appearance of cl-NOTCH1was absent in N1KO or N1N2KO after stimulation with JAG1 for 16 h. **B.** N1KO or N2KO partially blocked JAG1-induced inhibition of colony formation, while double N1K2 KO completely prevented growth inhibition. **C.** Infection with ectopic NFL1 cDNA restored NOTCH1 signaling in N1N2 KO cells. **D.** Re-Expression of NFL1 in PJ34 after N1N2KO restored JAG1-mediated inhibition of colony growth. **E.** Loss of both *NOTCH1* and *NOTCH2* (N1N2KO) in PJ34 protected cells from JAG1-induced morphology changes while re-expression of NFL1 restored spheroid formation caused by growth on JAG1.

**Supplementary Figure 4.**
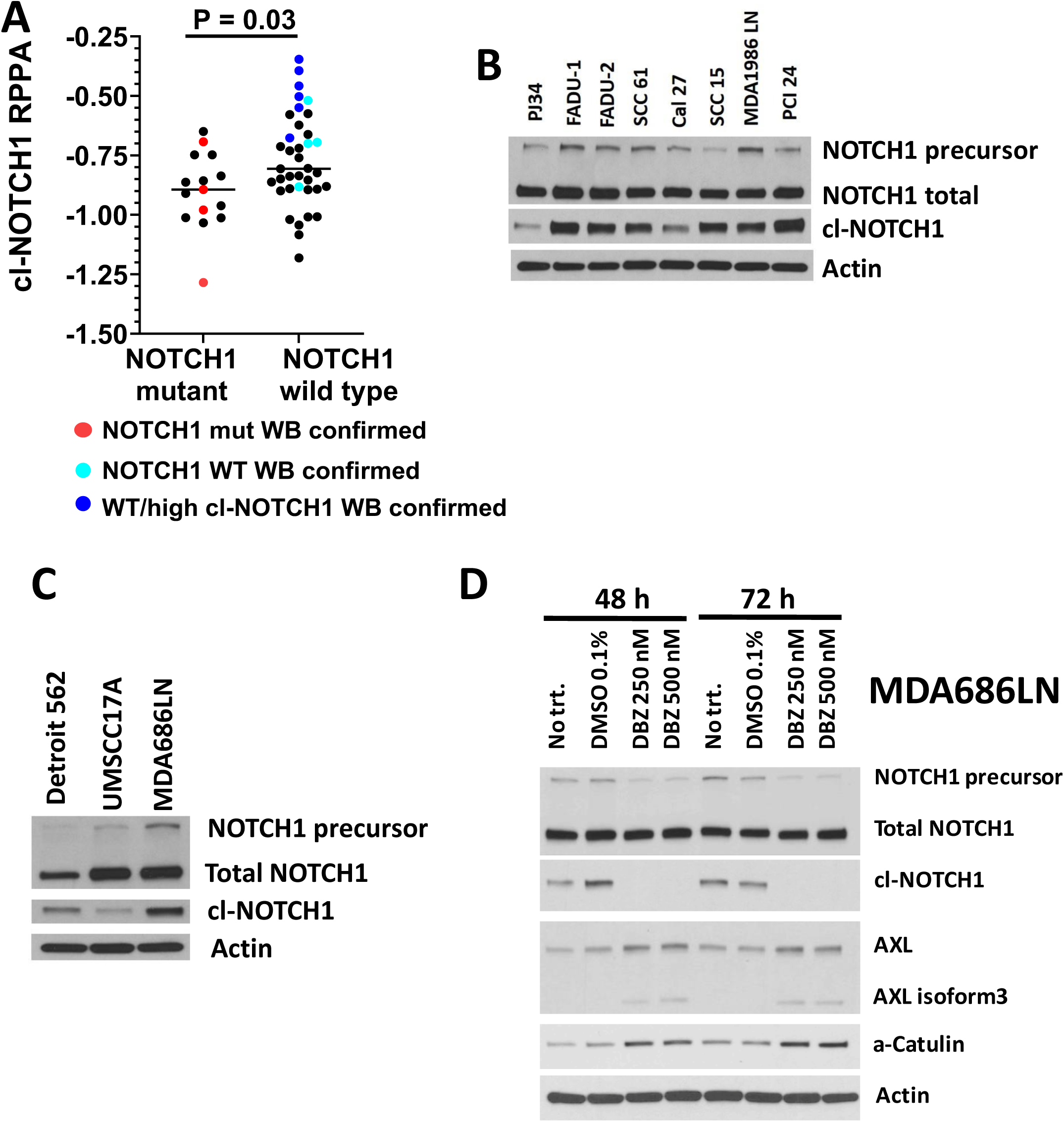
RPPA identifies *NOTCH1* WT cell lines with elevated baseline cl-NOTCH1expression which was inversely correlated with protein expression of AXL and a-CATULIN. **A.** Normalized RPPA values of cl-NOTCH1measured from *NOTCH1* mutant and *NOTCH1* WT HNSCC cell lines. Cell lines where cl-NOTCH1 protein was validated by western blots (WB) are indicated by colored symbols, with red dots for NOTCH1 mutants (HN31, UMSCC22A, UMSCC47, and HN4), light blue dots for NOTCH1 WT cells with low to moderate baseline cl-NOTCH (PJ34, 183, CAL27, and UMSCC1), and dark blue dots corresponding to *NOTCH1* WT cells with high basal cl-NOTCH1 (FaDu, PCI24, SCC61, SCC15, and MDA1986LN). **B.** Western blot confirmation of high cl-NOTCH1protein in subsets of WT cell lines, compared to PJ34 with low baseline activation. C. Confirmation of high baseline NOTCH1 activation in MDA686LN. **D**. DBZ inhibited cl-NOTCH1 protein which was associated with increased levels of both AXL and a-CATULIN protein.

**Supplementary Figure 5.**
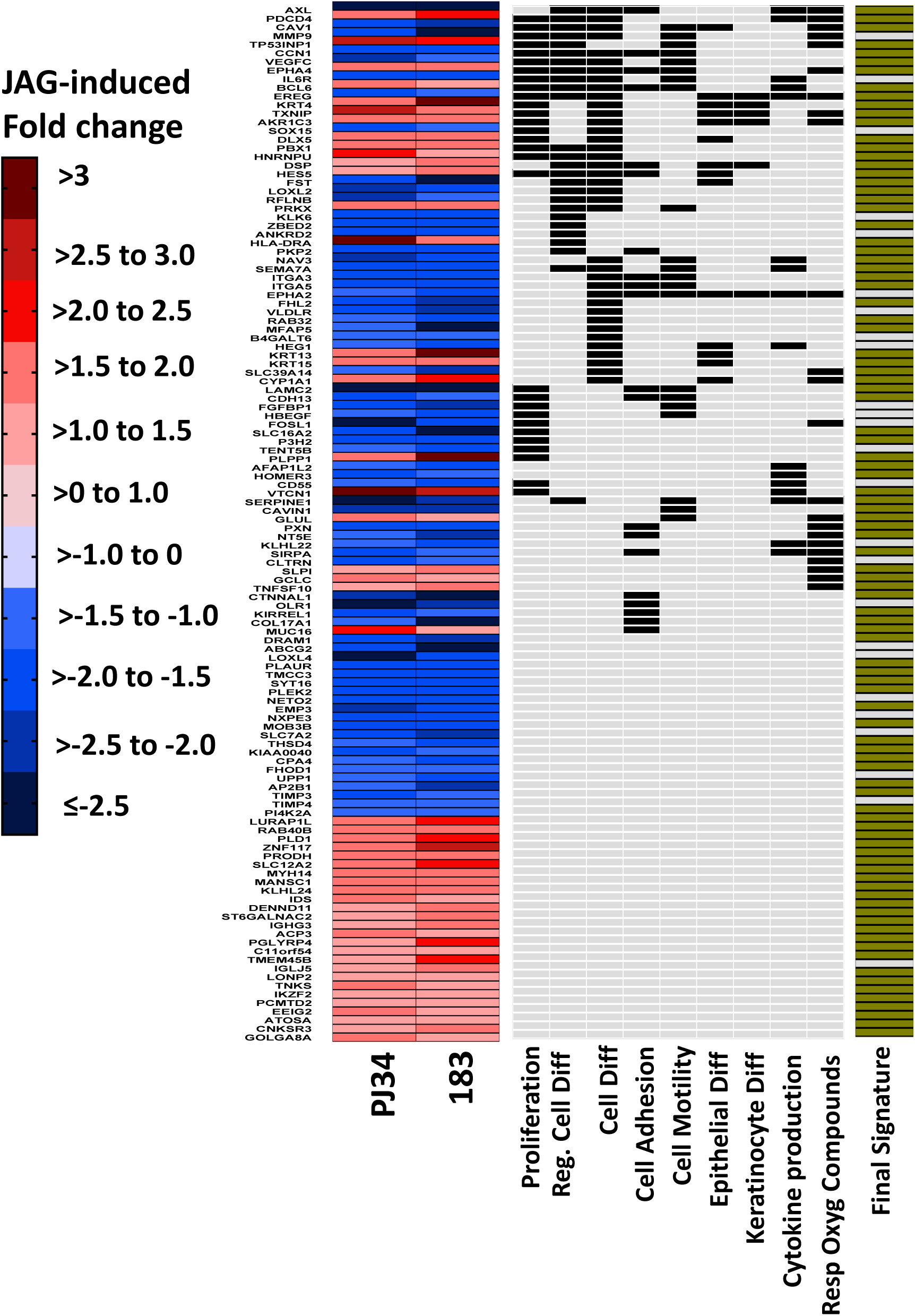
NOTCH1 regulates genes involved in proliferation, differentiation, attachment, motility, and response to oxygen containing compounds. The top 120 significant genes (i.e., minimum of 1.4-fold change) from *NOTCH1* WT PJ34 and 183 cells regulated in common by growth on JAG1 and their biological pathways. The fold change in each cell line after growth for 5 days on JAG1 is annotated vertically as a color gradient with red boxes representing increases, and blue boxes representing decreases. Membership in select GO pathways that were enriched in the gene set is annotated with black boxes, and gold boxes indicate genes that were kept for the final NOTCH1 gene activation signature discussed in the text.

**Supplementary Figure 6.**
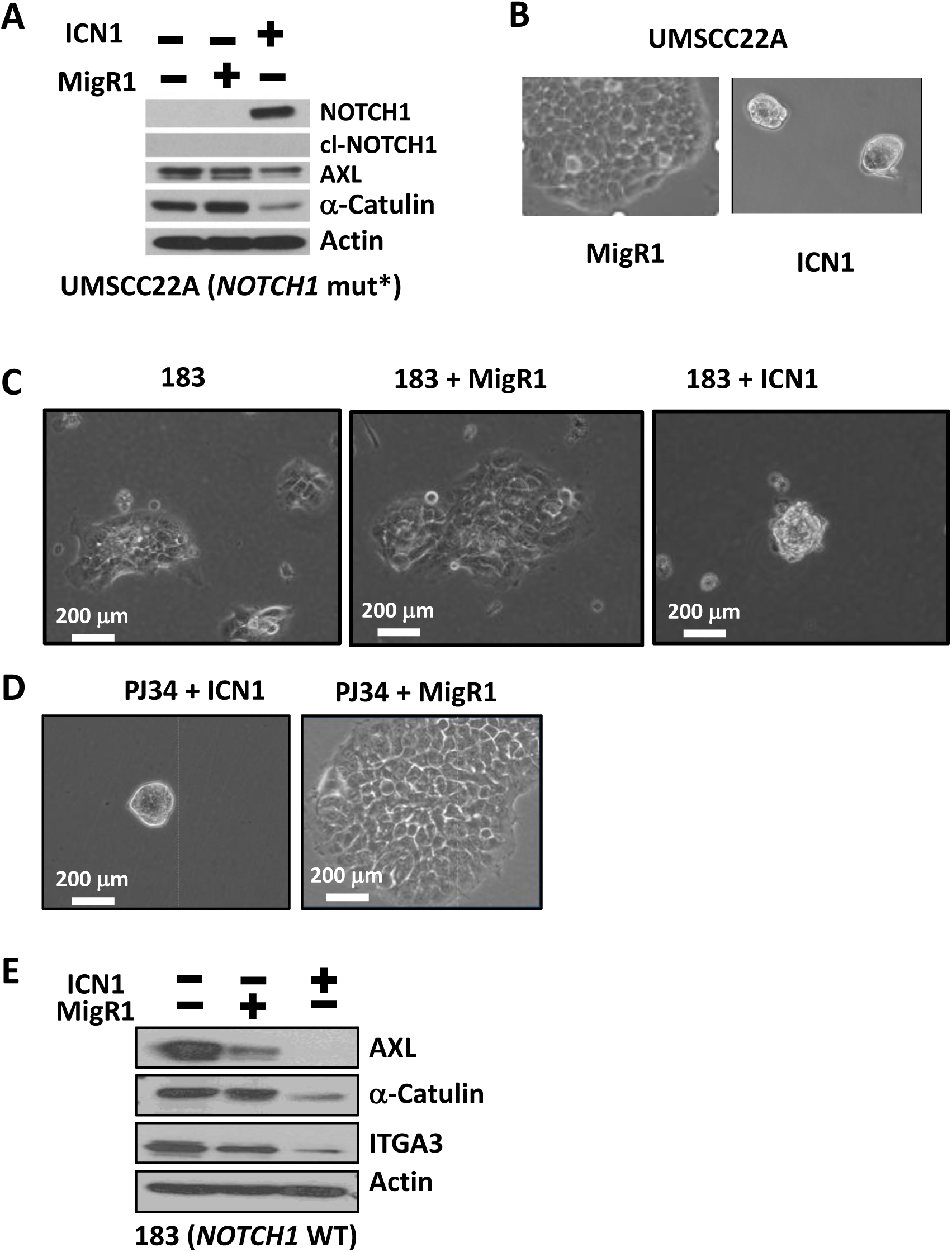
Ectopic expression of ICN1 mimics changes induced by JAG1 stimulation. **A.** Infection with ICN1 but not empty vector (MigR1) strongly inhibited protein expression of AXL and α-CATULIN. Widely used ICN1 cDNA product which begins several amino acids downstream of the native cleavage site is recognized by antibodies to the C-terminal region of NOTCH1 but not antibodies specific to the cleavage site. **B.** Expression of ICN1 in NOTCH1 mutant UMSCC22A produced the same morphological changes observed earlier upon expression of NFL1 and growth on JAG1. **C.** Expression of ICN1 in *NOTCH1* WT 183 produced the same morphological changes observed earlier with growth on JAG1. **D.** Expression of ICN1 but not MigR1 control in *NOTCH1* WT PJ34 produced the same morphological changes observed earlier with growth on JAG1. E. ICN1 expression triggered decreased protein expression of AXL, α-CATULIN, and ITGA3 in 183 cells.

**Supplementary Figure 7.**
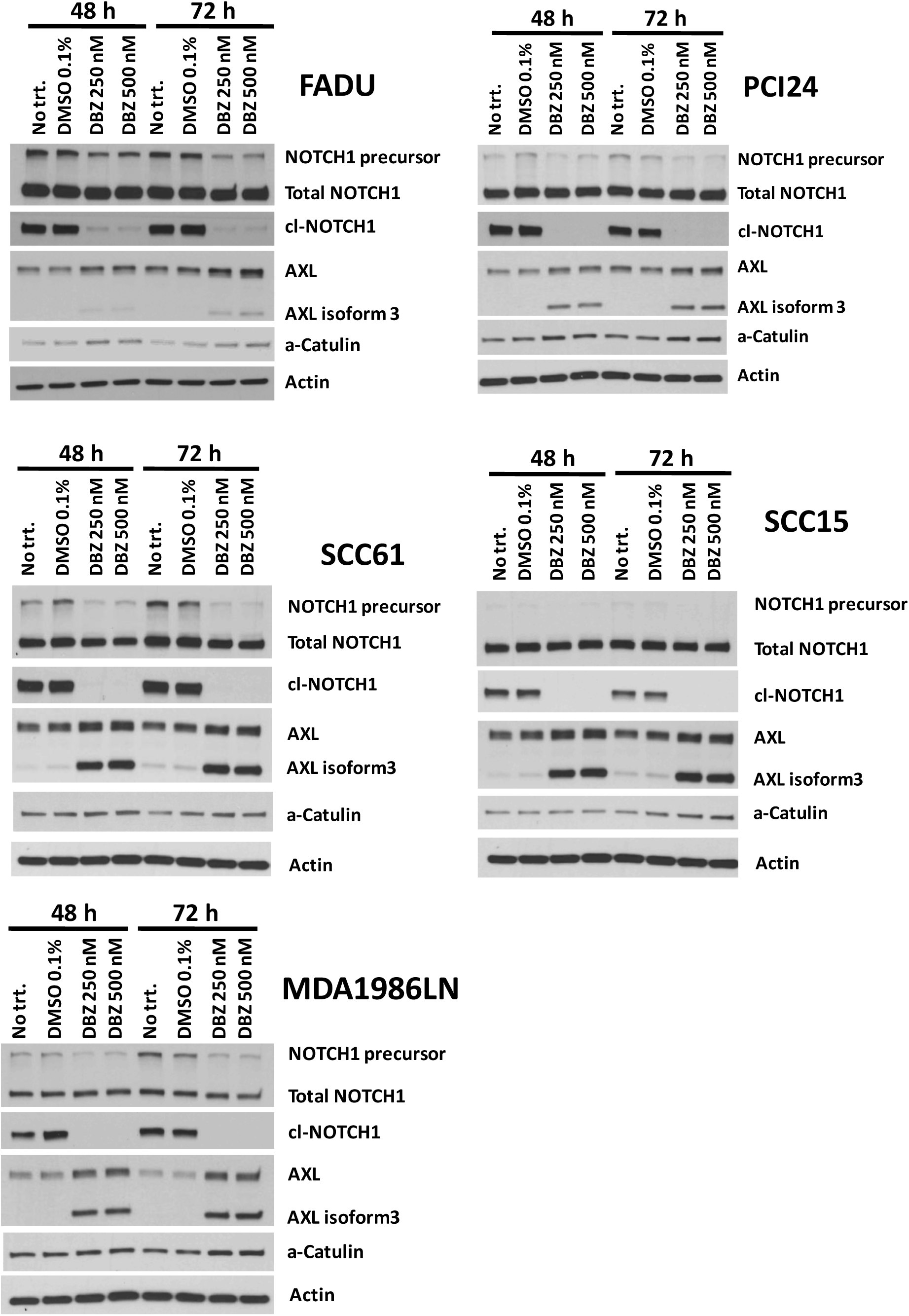
Evidence of NOTCH1 signaling in a subset of HNSCC tumor cell lines. Protein levels of both AXL and α-CATULIN increase when activated cl-NOTCH1 formation is blocked with the NOTCH inhibitor DBZ in five different *NOTCH1* WT cell lines with high endogenous NOTCH1 activity.

**Supplementary Figure 8.**
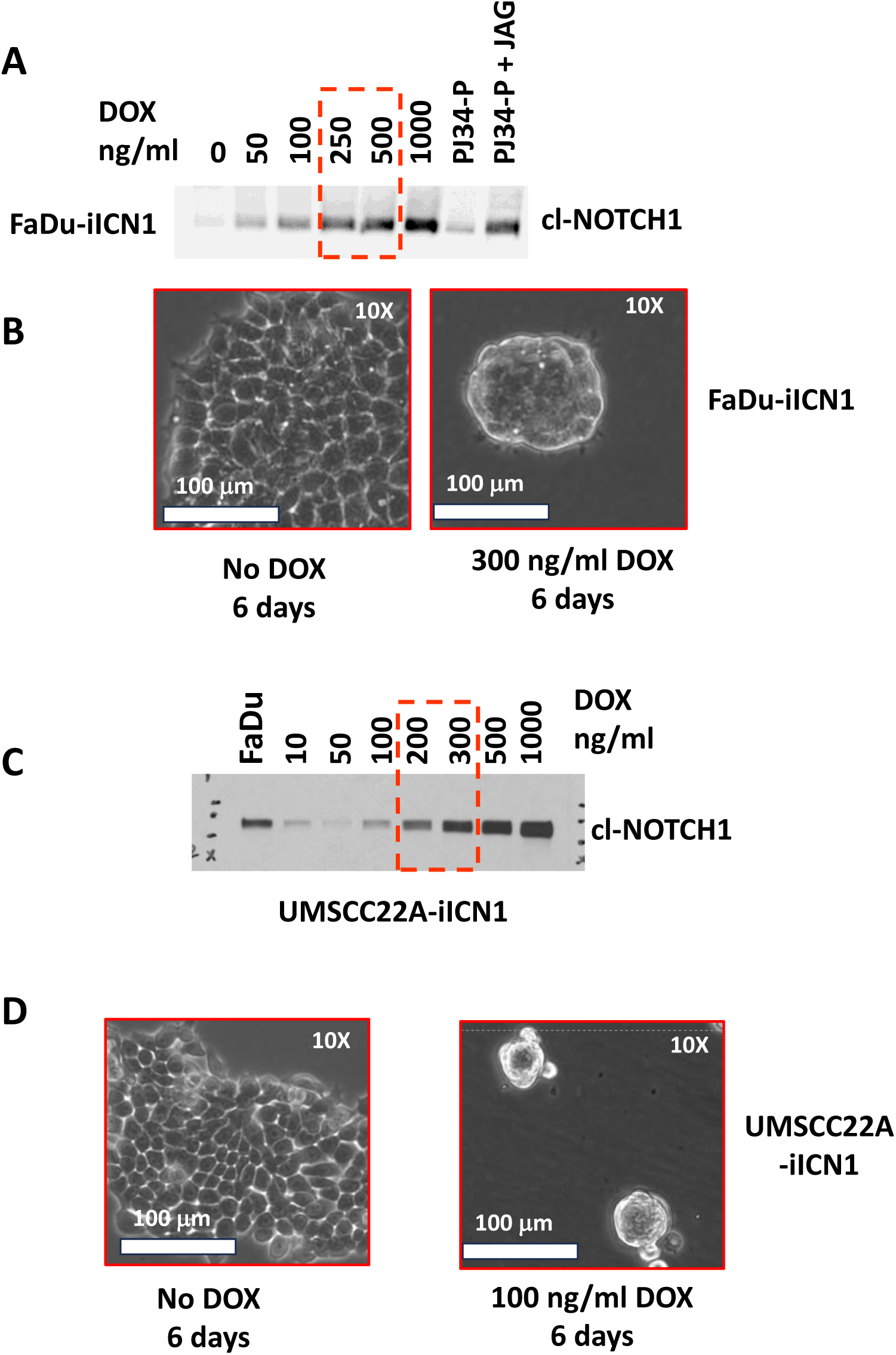
Physiological levels of iICN1 trigger morphology changes in *NOTCH1* WT and mutant tumor cell lines. **A.** Titration demonstrates that treating FaDu-iICN1 with 250ng-500 ng/ml DOX for 36 h induces protein levels of iICN1 equivalent to stimulating PJ34 with JAG1 for 16 h. **B**. As little as 300 ng/ml DOX for 6 days causes tumor spheroid formation in FaDu-iICN1. **C.** cl-NOTCH 1protein levels in UMSCC22A-iICN1 treated with 200-300 ng/ml DOX for 36 h are similar to basal levels in FaDu. **D.** Tumor spheroid formation in UMSCC22A-iICN1 cells after 6 days treatment with 100 ng/ml DOX.

**Supplementary Figure 9.**
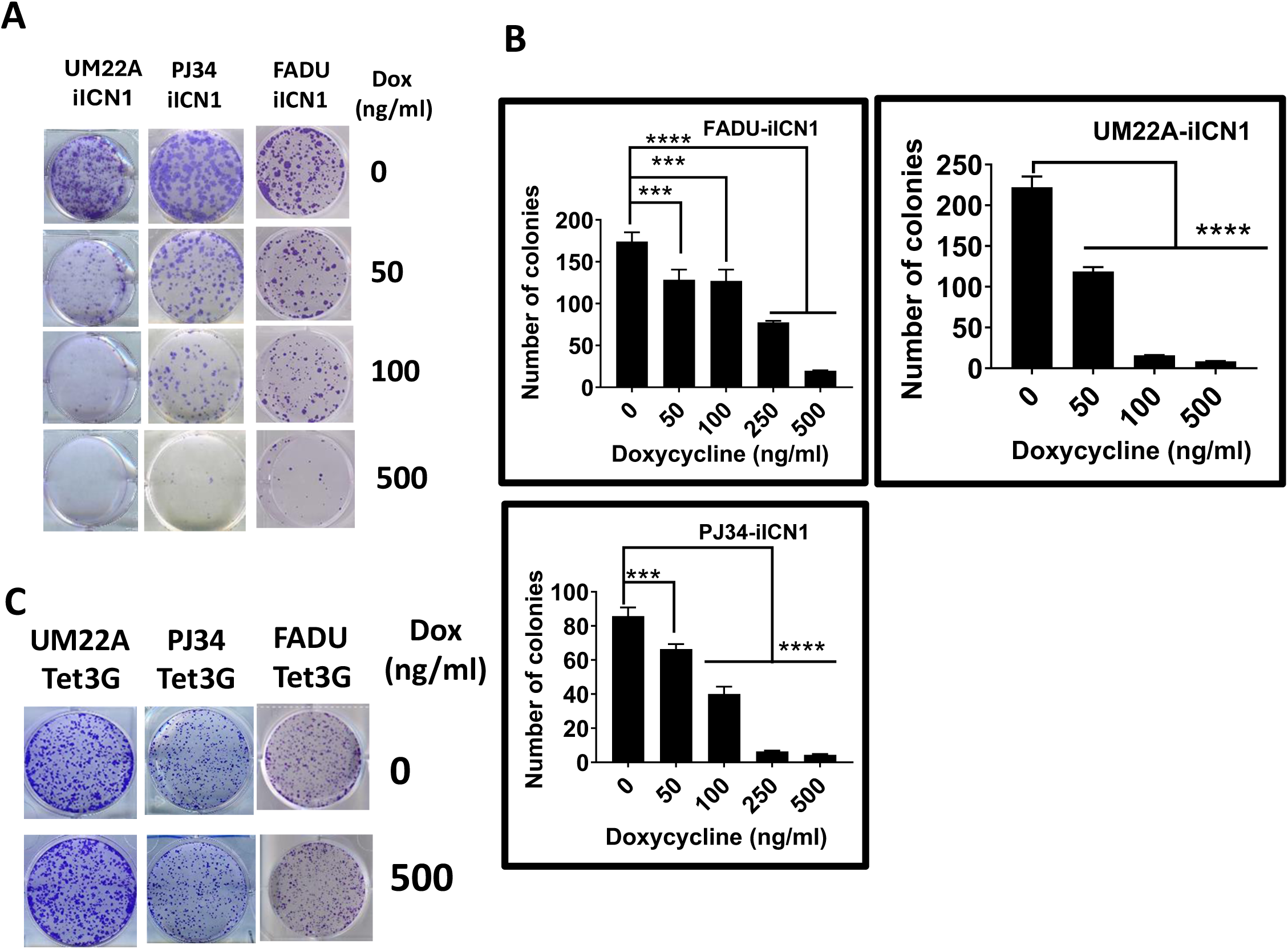
Dose response inhibition of colony formation by iICN1 in NOTCH1 mutant and WT cell lines. **A.** Crystal violet stain of colonies after increasing doses of DOX. **B**. Quantitation of colony inhibition after treating cells with increasing DOX doses. **C.** Treatment of control Tet3G expressing cell lines, lacking an iICN1 construct, with 1000 μg/ml DOX did not inhibit colony growth.

**Supplementary Figure 10.**
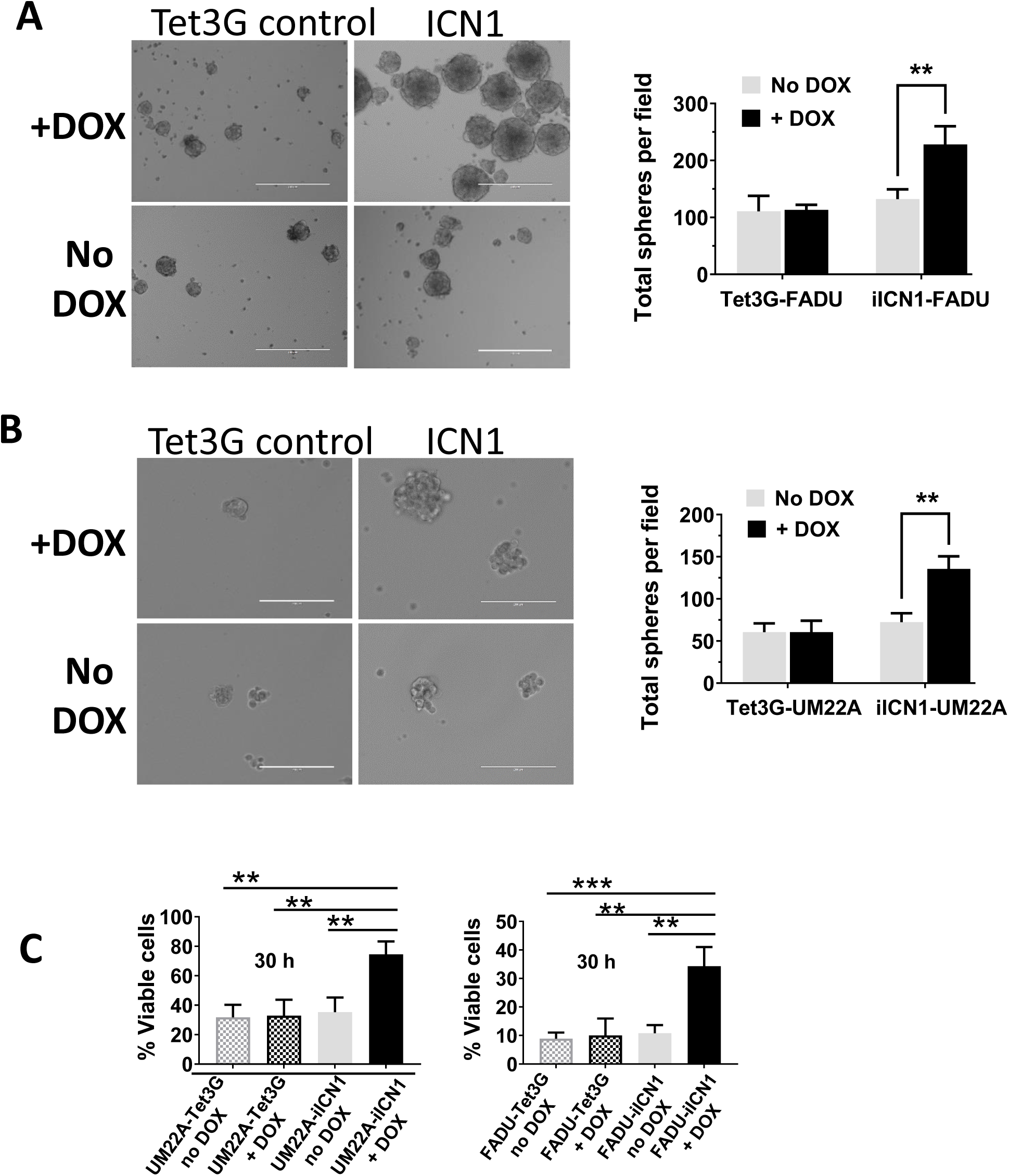
NOTCH1 activation increases tumor spheroid growth and anoikis resistance. **A.** Addition of 1000 mg/ml DOX increases the number of orospheres derived from FaDu-iICN1 in non-adherent low serum concentrations for 1 week. **B**. Increased orosphere numbers after growing UMSCC22A-iICN1 in 300 mg/ml DOX non-adherent low serum cultures for 1 week. **C.** DOX was added to attached UMSCC22A-iCN1 at 300 mg/ml, FaDU-iICN1 at 1000 mg/ml, or control Tet3G cells for 48 h before single cell suspensions were prepared in low serum containing media and an rotated in non-adherent tubes inside a humidified 37 ^0^C cell incubator for an additional 30 hours, in continued presence or absence of DOX. Viability of replicate cultures was examined with an automated cell counter.

**Supplementary Figure 11.**
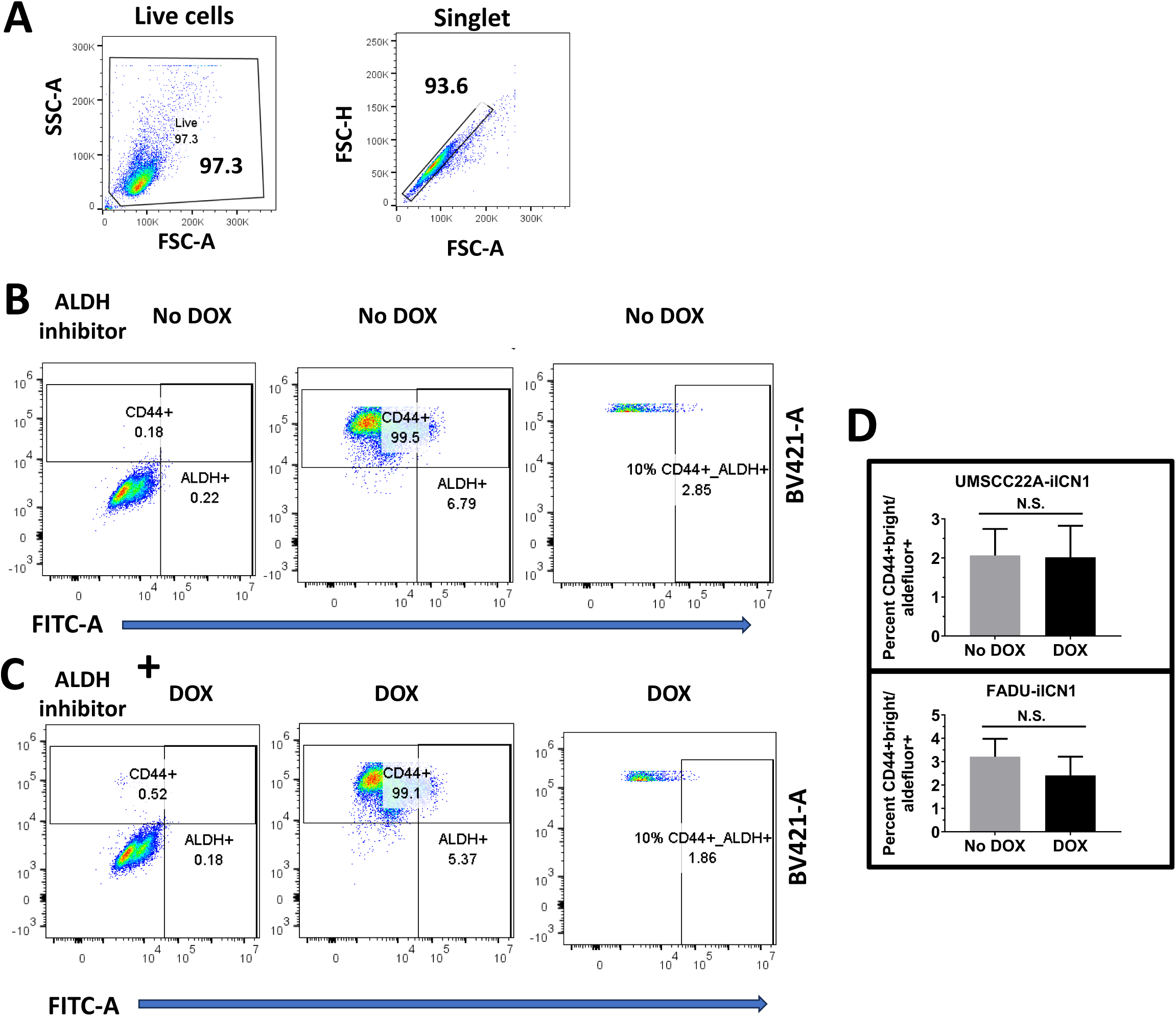
Activation of NOTCH1 does not increase the percentage of CD44+bright ALDEFLUOR-positive cells in NOTCH1 mutant or WT tumor lines. **A.** Flow cytometry gating strategy to identify live single cells. **B.** Representative gating of UMSCC22A-iICN1 cells in absence of Dox. The sample is split and one tube treated with a BV-421 conjugated isotype control antibody and the aldehyde dehydrogenase inhibitor (DEAB) before adding fluorescein conjugated substrate to set the background gates (left panel). The second tube is incubated with BV421-A conjugated anti-CD44 and Aldefluor substrate without *DEAB* to gate on the CD44+/Aldefulor+ population (middle panel) and a single fixed threshold is set for the top 10% brightest CD44+ (e.g., CD44+ bright), which is uniformly applied to all remaining UMSCC22A-iICN1 samples. The CD44+ bright population is further analyzed for the percentage of Aldefluor+ cells (right panel). **C.** A UMSCC22A-iICN1 sample pretreated with 200 ng/ml DOX for 48 h to induce ICN1 before staining is gated in the same fashion, except Aldefluor gating is set with a sample specific tube treated with DEAB (left panel) and CD44+bright cells are analyzed in a companion tube without DEAB for percentage of Aldefulor+ using the previously set CD44+ threshold (right panel). D. The total percent of CD44+bright/Aldefluor+ cells does not increase after iICN1 induction in either UMSCC22A-iICN1 or FaDu-iICN1.

**Supplementary Figure 12.**
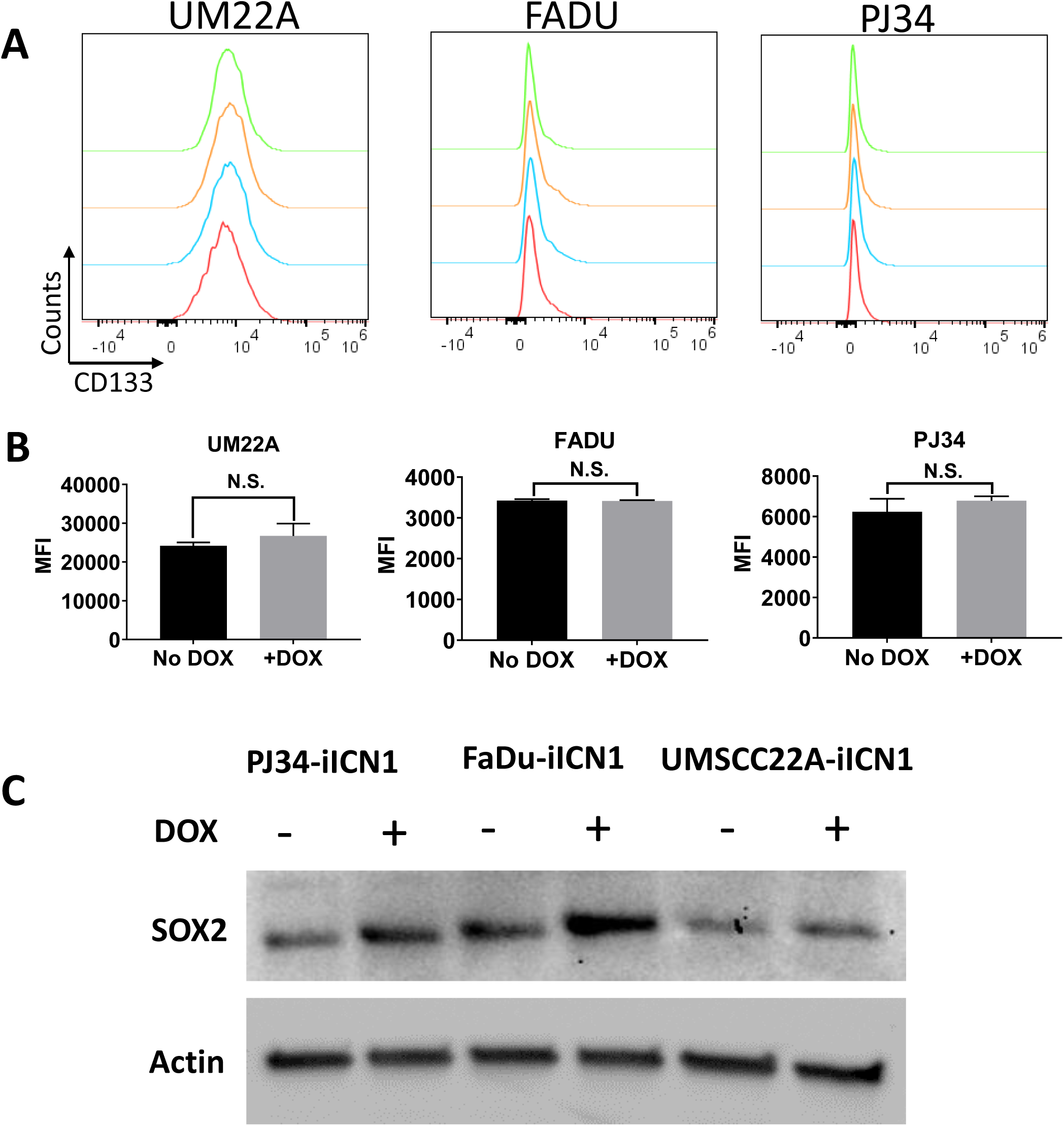
Activation of NOTCH1 fails to increase surface CD133+ but increases expression of SOX2 in *NOTCH1* mutant (UMSCC22A-iICN1), or *NOTCH1* WT cell lines (FaDu-iICN1 and PJ34-iICN1). **A.** Cells were incubated for 48 h with or without DOX at 200 ng/ml (UMSCC22A), 300 ng/ml (FaDu), or 1000 ng/ml (PJ34) before staining with antibody to surface CD133 by flow cytometry. Fluorescent intensity histograms are shown for control biological replicates without DOX (red and light blue traces) or after DOX treatment (green and orange traces). **B.** Statistical comparison of CD133 mean fluorescence intensity (MFI). **C.** Western blot analysis of SOX2 protein expression in similarly treated cells.

**Supplementary Figure 13.**
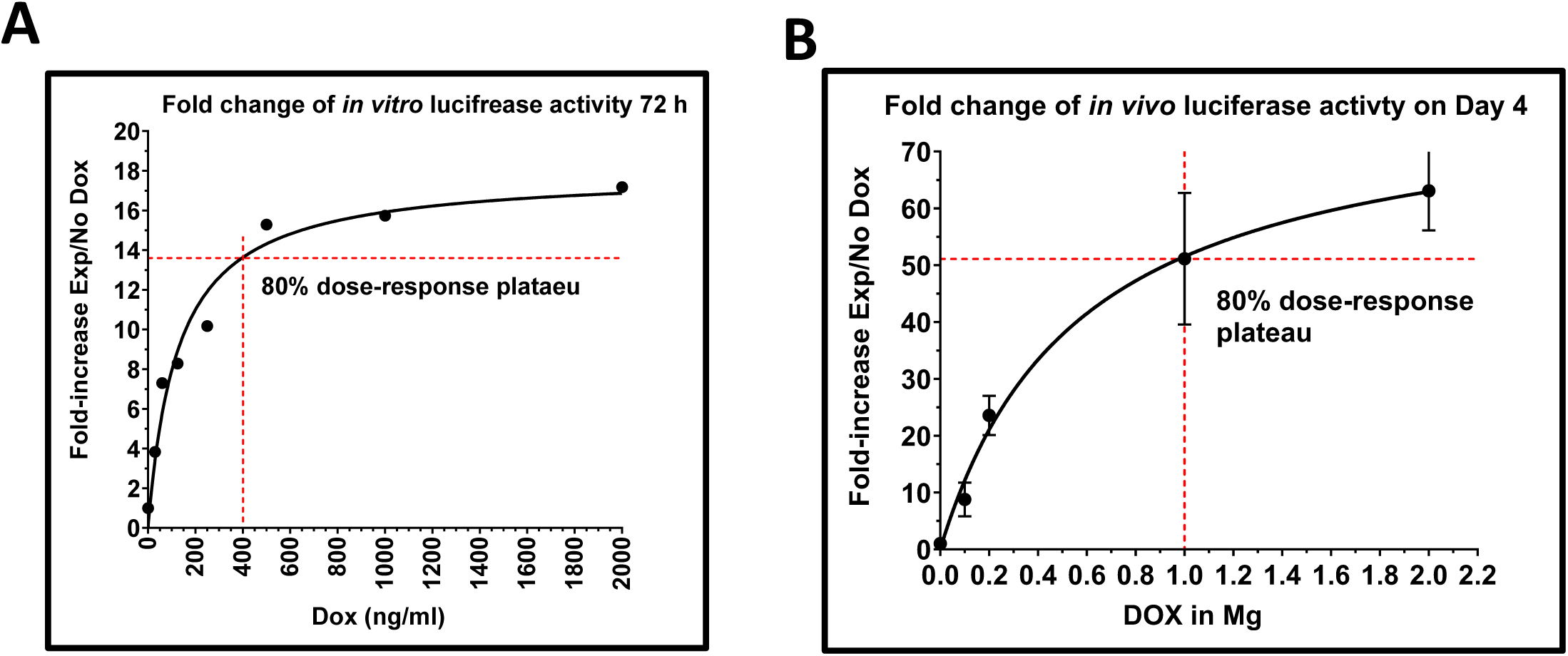
*In vitro* and *in vivo* dose response of the inducible DOX promoter. **A.** UMSCC22A Tet3G reporter cells stably expressing firefly luciferase cloned into pLVX-TRE3G-mCherry were seeded into 6 well plates and luciferase activity measure in a plate reader 36 hours after induction with increasing concentrations of DOX. A dose of ~400 ng/ml resulted in luciferase activity that was 80% of the *dose response* plateau, but not outside linear response of the plate reader **B**. Standard curve relating relative *in vivo* luciferase luminescence to DOX concentration given to mice by oral gavage for 4 days, demonstrating that 1 mg DOX yielded activity that was 80% of the in *vivo* plateau.

**Supplementary Figure 14.**
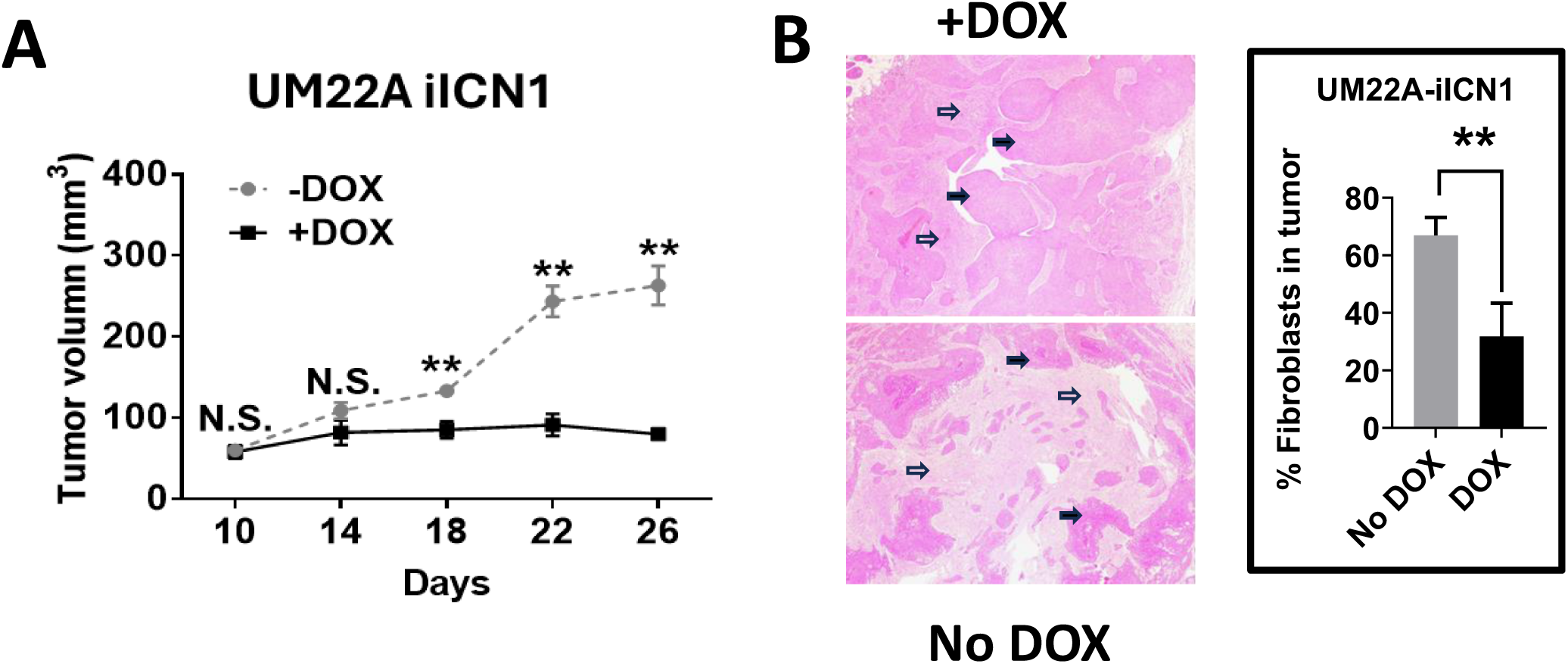
Activation of NOTCH1 signaling in *NOTCH1* mutant tumors profoundly inhibits *vivo* tumor growth and reduces the number of CAF cells. **A.** UMSCC22A-iICN1 was inoculated into mouse flanks and mice were randomly assigned to receive either no DOX or daily DOX (1 mg) by oral gavage for two weeks. Tumor volumes were compared over time. **B**. H&E staining of a representative tumor that eventually formed in mice treated with DOX (top panel) and a tumor that formed in the absence of DO X (bottom panel). Filled arrows indicate tumor nests, while hallow arrows designated areas predominately populated by CAFs. The percentage of CAFS in tumors was significantly lower when ICN1 was induced with DOX.

**Supplementary Figure 15.**
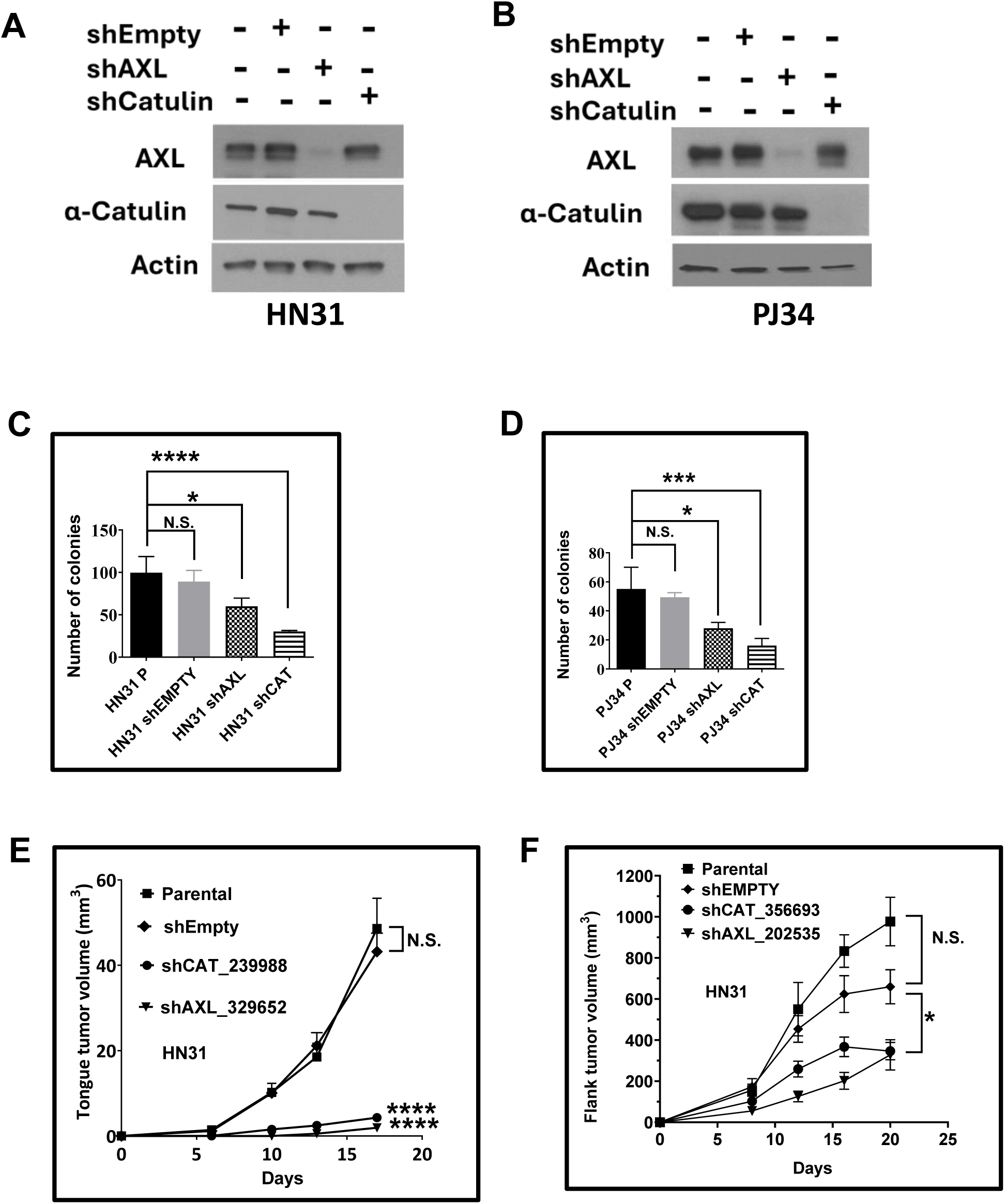
Knockdown of *AXL* and a-CATULIN to mirror NOTCH1 signaling inhibits *in vitro* and *in vivo* tumor growth. **A.** Western blot confirmation of AXL and a-CATULIN protein inhibition following shRNA KD in *NOTCH1* mutant HN31 cells 96 h post-infection with specific shRNA. **B.** Western blot confirmation of AXL and a-CATULIN protein inhibition following shRNA KD in NOTCH1 WT PJ34 cells 96 h post-infection with specific shRNA. **C.** KD of AXL and a-CATULIN blocks in vitro colony formation in HN31. **D.** KD of AXL and a-CATULIN blocks *in vitro* colony formation in PJ34. **E.** KD of AXL and a-CATULLIN profoundly inhibits HN31 *in vivo* tumor growth in an orthotopic tongue tumor model. E. **E.** KD of AXL and a-CATULLIN with alternate shRNA vectors targeting different regions inhibits HN31 *in vivo* tumor growth in a subcutaneous flank model.

**Supplementary Figure 16.**
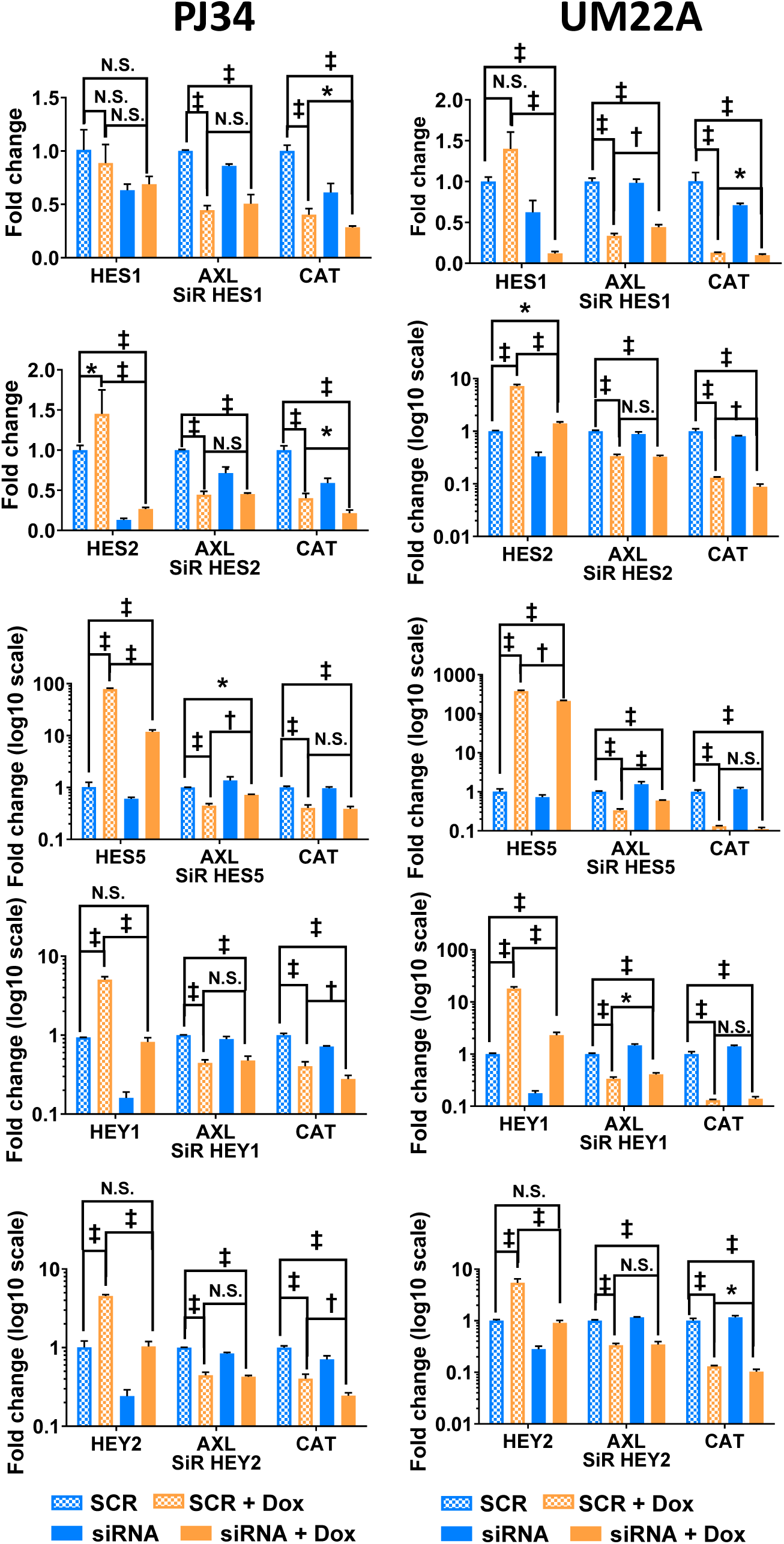
Canonical HES/HEY family members are not key mediators of ICN1 induced reductions in *AXL* and a-*CATULIN* expression. PJ34-iICN1 (left) or UMSCC22A-iICN1 (right) were pretreated with siRNA to HES1, HES2, HES5, HEY1, or HEY2 or control non-targeting siRNA (SCR) for 48 h before replating in the absence or presence of 300 ng/ml DOX for 24 h. Expression of the HES/HEY knockdown targets, AXL, and α-Catulin were quantitated by qPCR. * P <0.05, ^†^ P < 0.001, ^‡^ P <0.0001.

**Supplementary Figure 17.**
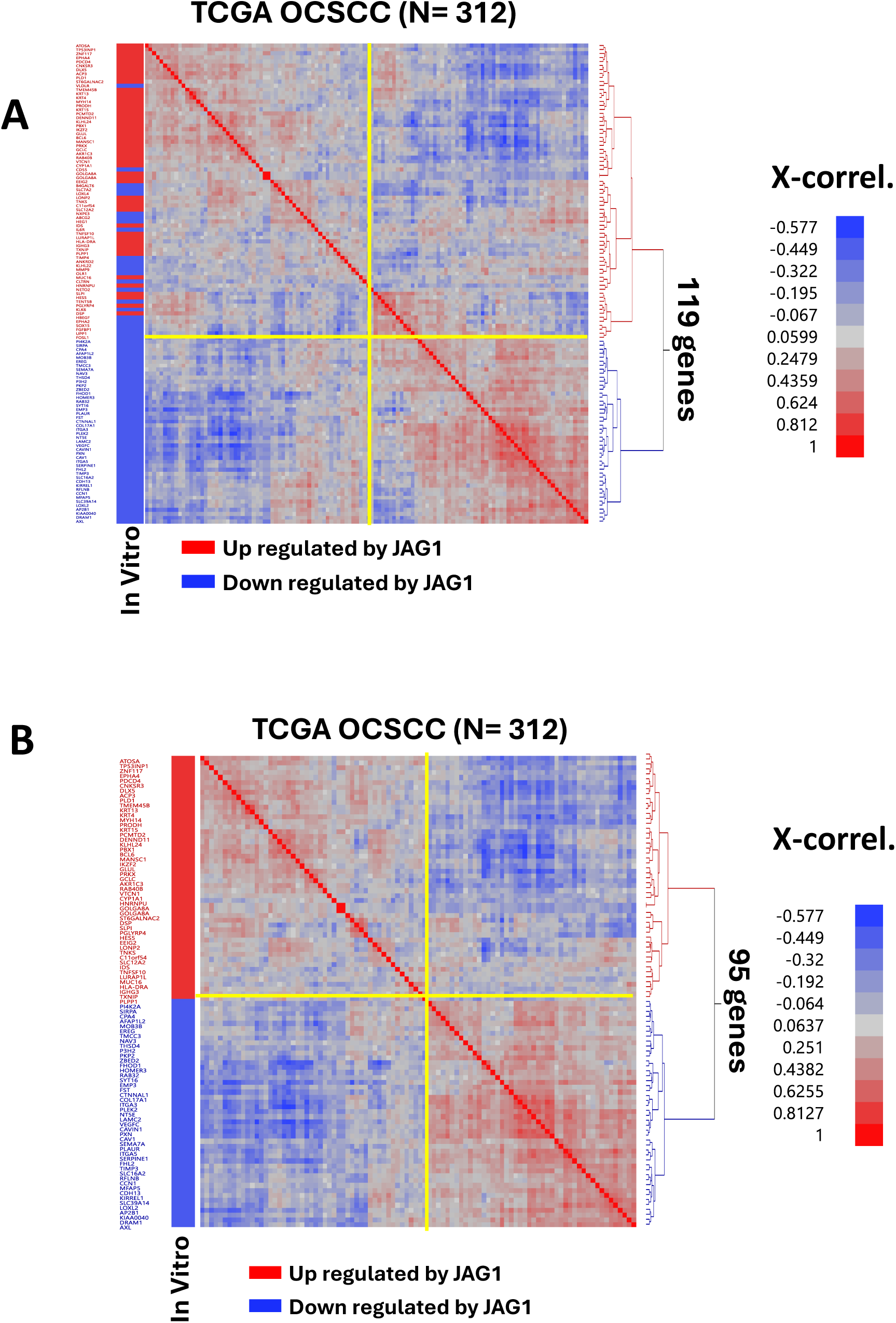
Development of an in vivo NOTCH1 signaling gene expression signature. **A.** The TCGA OCSCC RNA-seq cohort was mined for cross-correlation of RNA expression using gene candidates from the list of 119 top genes regulated *in vitro* after JAG1 exposure. Hierarchical two-way clustering of cross-correlation coefficients identified two main gene clusters annotated by whether the genes were up-regulated (red box) or down regulated (blue box) after JAG1 exposure. **B.** Re-clustering TCGA OCSCC cross-correlation coefficients with the remaining 95 genes after removal of those with inconsistent *in vitro* and *in vivo* behavior.

**Supplementary Figure 18.**
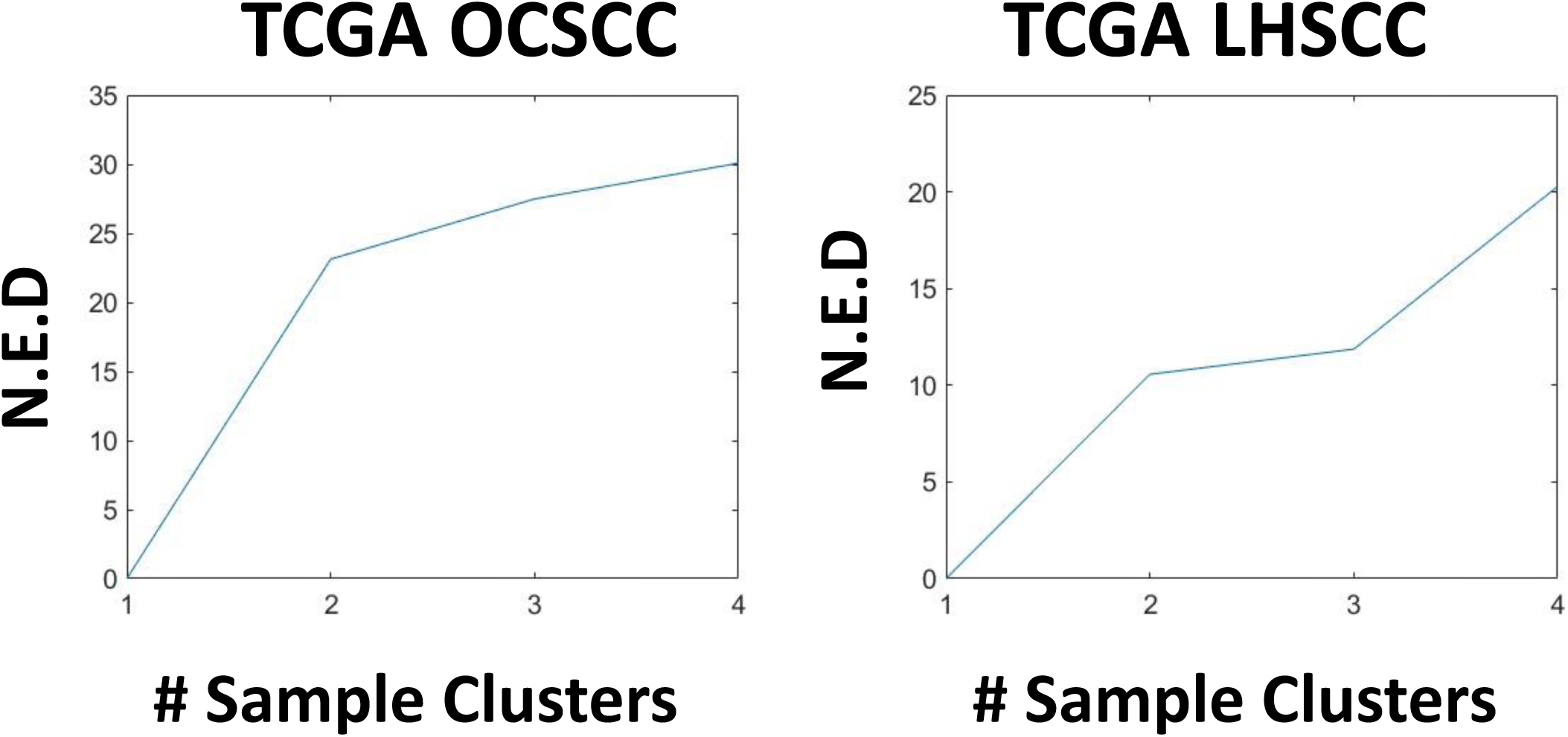
Selection of optimal sample clusters. Consensus hierarchical clustering of TCGA OCSCC and LHSCC samples using the 95 gene NOTCH1 signature was performed and the similarity matrices achieved using increasing numbers of sample clusters (e.g. N=1 to 4) were compared to theoretical perfection matrices to select a local minimum for the normalized Euclidean distances (N.E.D.) to identify an optimal number of sample clusters, which happened to be N =2 clusters for each disease subsite.

**Supplementary Figure 19.**
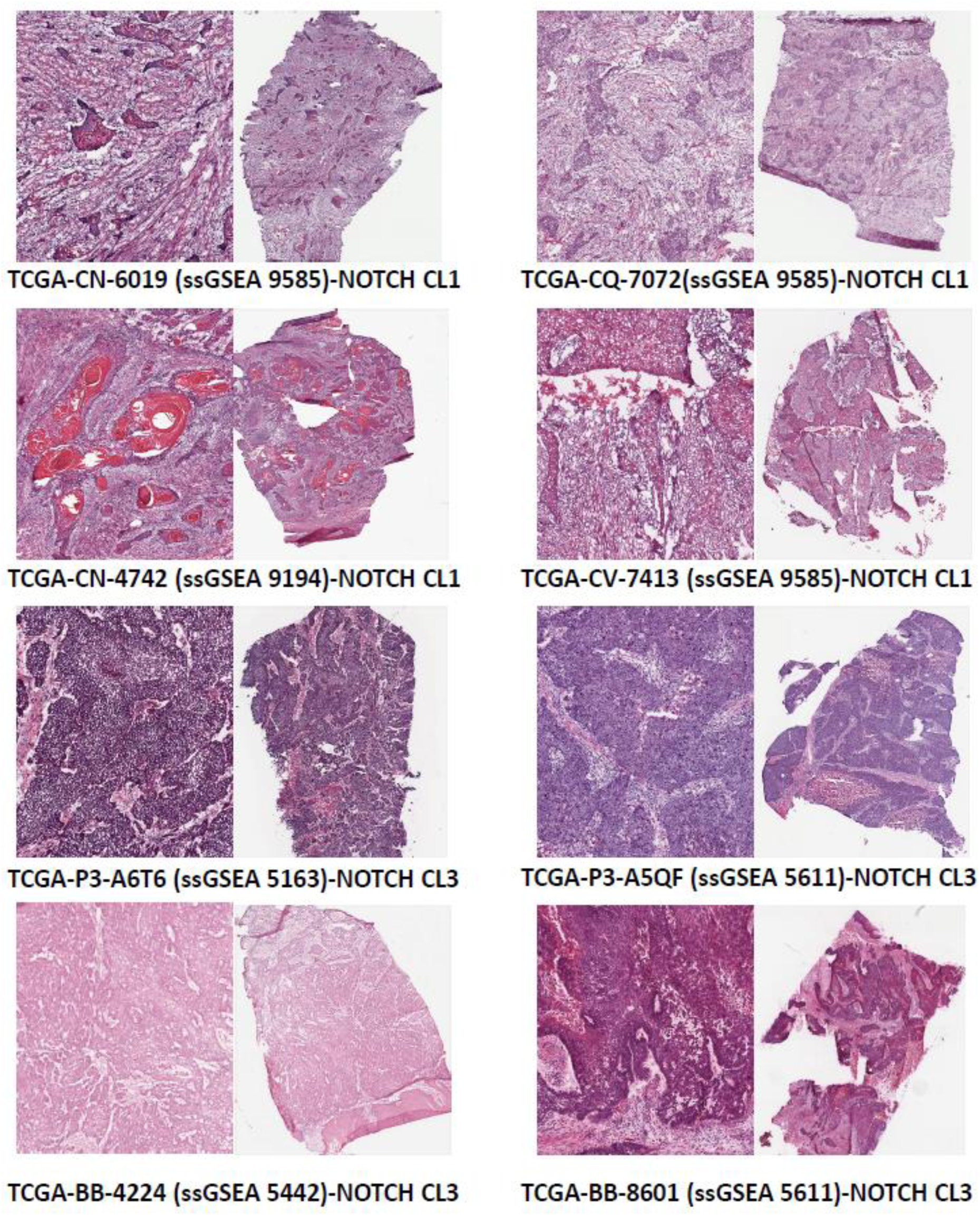
Visual confirmation that TCGA tumor samples identified with active NOTCH1 signaling contain fewer CAF. H&E images were downloaded from the TCGA data portal and the CAF ssGSEA scores appear in parenthesis alongside the sample TCGA identification numbers.

**Supplementary Figure 20.**
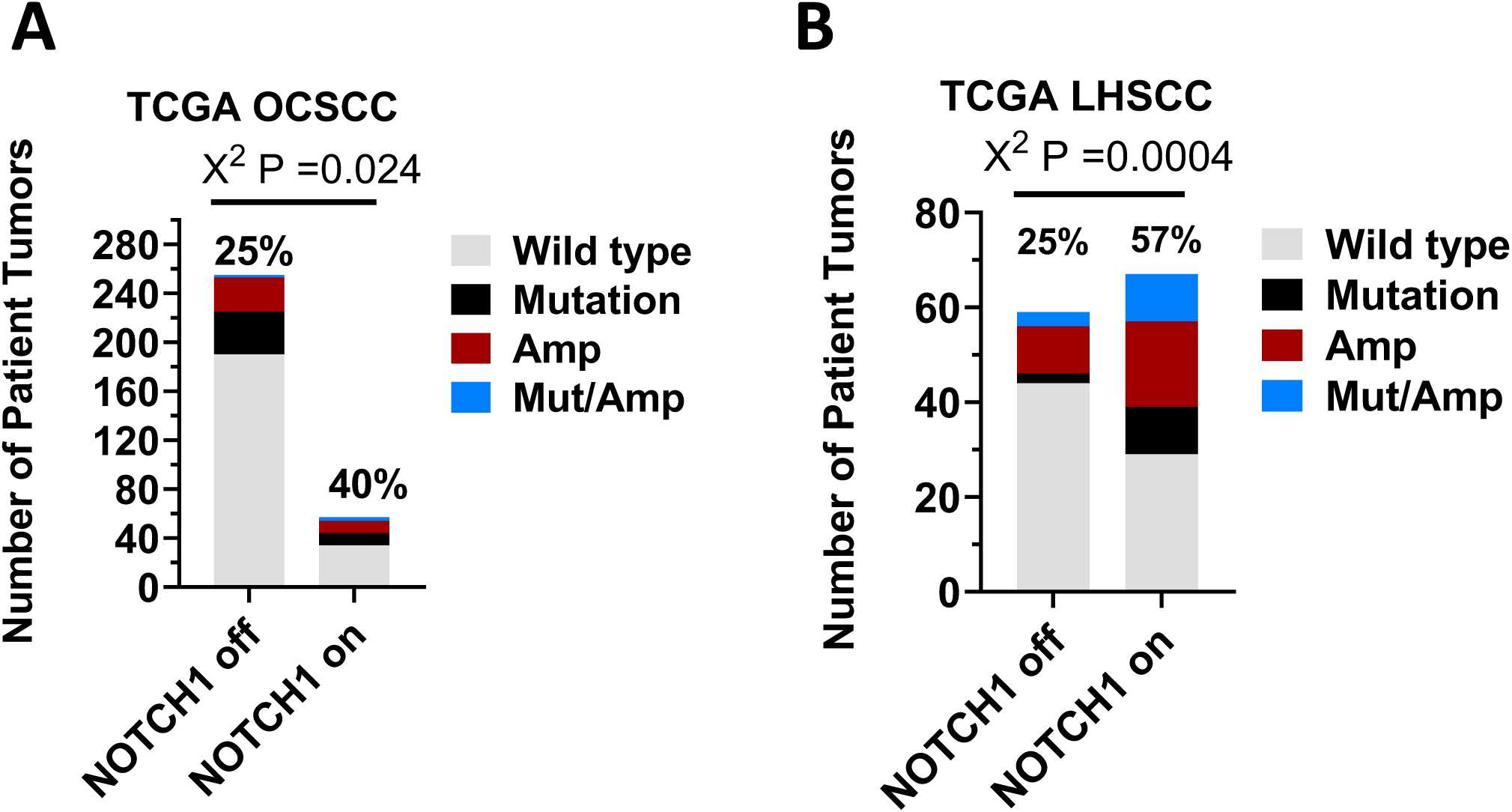
Patients lacking NOTCH1 mutations are significantly enriched for genomic alterations in PIK3CA. The proportion of tumors with genomic alterations in PIK3CA, including mutations and/or high-level gene copy gains, is significantly higher in in the group associated with NOTCH1 signaling identified through hierarchical clustering of TCGA OCSCC (A) or (B) LHSCC.

## Funding to acknowledge

This work was supported by the National Institute of Dental and Craniofacial Research through RO1DE024179 (M.J.F.), the National Cancer Institute (NCI) through RO1CA235620 (M.J.F.), U01DE025181 (V.C.S), and the Cancer Prevention Institute of Texas through RP200369.

