## supplementary methods for "NOTCH1 Acts as a Tumor Suppressor That Induces Early Differentiation in Head and Neck Cancer"

**Viral vectors, shRNA Vectors, and siRNA Reagents.**

A 7.3 kb cDNA encoding WT full length human NOTCH1 receptor (NFL1) with a Kozak sequence was provided by Origene and digested with EcoR1/XhoI to subclone into an empty retroviral vector MigR1 (Addgene), where the multiple cloning site (MCS) that had been previously modified to contain MfeI (EcoR1 compatible) and Xho1 sites from 5’ to 3’. Sanger sequencing confirmed that the entire NFL1 cDNA insert matched the reference sequence for full length human NOTCH1 (NM 017617) encoding 2555 amino acids. The MCS of MigR1 is upstream of an IRES-EGFP (enhanced green fluorescence protein) cassette so NFL1 cDNA is expressed from the same mRNA as EGFP, allowing purification of infected cells by flow cytometry. An ICN1 construct encoding human NOTCH1 Pro1770 to Lys2555 cloned into MigR1 was obtained from Dr. Patrick Zweidler-McKay. To express inducible activated cleaved NOTCH1 that could be recognized by cl-NOTCH1 antibodies we utilized RNA from NOTCH1 WT FaDu to generate a cDNA fragment encoding Val1754 to Lys2555 which was amplified with primers containing a Kozak sequence and artificial ATG methionine start site. Amplified cl-NOTCH1 cDNA was then cloned into the tetracycline/doxycycline lentiviral plasmid pLVX-TRE3G-mCherry using an In-Fusion cloning kit (Takara). GIPZ shRNAs used for targeting AXL and a-CATULIN (CTNNAL1) were from Horizon and included, V2LHS_239988 (AXL, AACTTAGATGCTTAGGATC), V3LHS-329652 (AXL, TGAGGATGGAGTCGTCCTG), V2LHS-239988 (CTNNAL1, TATTTCCAGAGGTTCTGTC), and V3LHS_356693 (CTNNAL1, TGTTTTCTTGCTTTGAGCT). SMARTPool siRNA mixtures targeting either human HES1, HES2, HES5, HEY1, and HEY2 were purchased from Horizon.

CRISPR-Cas9 NOTCH1 (sc-400167-KO-2), and NOTCH2 (sc-401323-KO-2) plasmids were obtained from Santa Cruz.

**Antibodies**

Antibodies to cl-NOTCH1 (#4147), AXL (#8661), SOX2 (#23064) and LAMC2 (#53884) were purchased from Cell Signaling Technology (Danvers, MA). Antibodies to total NOTCH1 (Sc 6014), total NOTCH2 (sc 5545), α-Catulin (sc 390584) were from Santa Cruz Biotechnology (Santa Cruz, CA). The β-Actin antibody (A1978) was bought from Sigma (St Louis, MO), and anti-ITGA3 (#PA5-143239) was purchased from ThermoFisher (Waltham, MA).

**Viral Mediated Ectopic Gene Expression and shRNA Knockdown (KD)**

Retroviral and lentiviral particles were generated by transfecting expression plasmids into helper cell lines—HEK-293GP2 for retroviruses and HEK-293F17 for lentiviruses. For retroviral production, a mixture of MigR1, ICN1, or NFL1 (10.8 mg) with 1.2 mg of the packaging plasmid pCMV-VSV-G (Addgene) was transfected into a 10 cm dish of 6 million adherent HEK-293GP2 cells using GenJet Plus (SignaGen) under a modified protocol that minimized volumes during the initial 45 minutes at 37 °C in a tissue culture incubator. After this period, complete media was added for an additional 8 hours at 37 °C before the transfection reagents were washed out. The cells were then incubated at 33 °C to optimize viral production, and the virus-containing supernatants were harvested at 48 and 72 hours, centrifuged, and pooled. Lentiviral particles (e.g., shRNA or inducible expression plasmids) were produced similarly, using a mixture of 10 µg expression plasmid, 5 µg packaging plasmid pCMV-dr8.2 (Addgene), and 5 µg envelope plasmid pMD2.G (Addgene) to transfect HEK-293F17 cells with GenJet Plus. The cells were incubated at 37 °C, and virus was harvested from the supernatants at 48 and 72 hours.

Cells were infected with retroviral or lentiviral particles using a reverse spin inoculation protocol. Briefly, trypsinized tumor cells were counted and mixed with serial dilutions of viral supernatant, fresh media, and polybrene (final concentration of 8 μg/mL). One million cells were then seeded in 2 mL per well in 6-well plates. The plates were centrifuged at 12,000 × g for 1.5 hours at room temperature, then transferred to a 37 °C tissue culture incubator for 48 hours. Infected cells were subsequently purified by flow cytometry or, in some cases, selected using antibiotics. Cell lines expressing DOX-inducible lentiviral constructs were generated by first infecting cells with pLVX-Tet3G (Clontech) encoding the Tet-ON 3G regulator protein and selecting for G418 resistance to obtain Tet3G modified cells. Cells stably expressing Tet3G were then infected with human ICN1 that had been subcloned into pLVX-TRE3G-mCherry (iICN1) and selected for 1 week in puromycin. For some experiments, polyclonal puromycin selected cells were used while in other cases, cell sorting was used to generate and screen clones with minimal leakiness and maximal ICN1 induction.

**Analysis of RNA expression.**

Total RNA was isolated from replicate samples of PJ34 or 184 cells grown for 5 days on plates coated with either JAG1 or control FC. The RNA isolated with Trizol was purified by ethanol precipitation and hybridized to Affymetrix HuGene 2.0 ST arrays by the MD Anderson Sequencing and Microarray Core Facility. Data was processed using aroma.affmetrix package in R to quantify the CEL files with Robust Multiarray Averages (RMA) , apply background correction, quantile normalization, and RMA Probe-level summarization. The processed data was log2 transformed before analysis of differentially expressed genes (DEGs) using a linear model fit with both treatment and cell line as fixed effects. P values were modeled using a beta-uniform mixture model and combined with false discovery rate at 0.05 to determine P value cutoffs.

RNA-seq was used to identify genes differentially expressed after iICN1 expression in PJ34. Replicate cultures of either PJ34-Tet3G or PJ34-iICN1 cells were incubated in the presence or absence of 1000 ng/ml DOX for 36 h and total RNA isolated with an RNeasy Kit was sequenced by the MD Anderson Sequencing and Microarray Core Facility. Gene expression was normalized as counts per million and log2 transformed before identifying differentially expressed genes using the response screening module in JMP13 statistical software, which conducts individual T-tests for every gene and applies a Benjamini-Hochberg correction (FDR = 0.1) to calculate adjusted P values. To minimize the number of tests, poorly expressed genes were filtered out before the analysis by removing any gene whose average for at least one treatment group failed to exceed a low expression threshold (e.g., log2 expression < 2). Data from control Tet3G cells were used to identify and exclude any genes regulated by DOX alone, in the absence of iICN1 expression.

**Consensus Hierarchical Agglomerative Clustering.**

Z scores from select genes were employed in a two‐way consensus hierarchical agglomerative clustering analysis using Ward’s minimum variance method, implemented via a custom Matlab script (available at <https://github.com/aif33/Hierarchical-two-way-agglomerative-consensus-clustering>) that we previously described (30). This approach is based on a modification of the resampling method described earlier by Monti et al., wherein 80% of the samples are randomly selected without replacement in each iteration. For each resampled set, Ward’s clustering partitions the samples into N clusters, with N varied over a user-specified range (e.g., 2, 3, 4, etc.), and the frequency with which any two samples co-occur in the same cluster is recorded in a similarity matrix. This matrix is then transformed by retaining the original similarity values for pairs that consistently cluster together and replacing the values for pairs that do not with one minus the similarity value, effectively representing the fraction of iterations in which the samples did not co-cluster. Ideally, this transformation would yield an identity matrix, with all off-diagonal values equal to one, indicating perfect separation. The deviation of the observed transformed matrix from this ideal is quantified by computing the Euclidean distance between the two matrices, which is then normalized by dividing by N to yield a normalized Euclidean distance (NED). The optimal number of clusters is determined by identifying a localized minimum on the NED versus cluster number plot, thereby balancing the tradeoff between increasing cluster granularity and the preservation of meaningful information. For graphical representation, the untransformed similarity matrix was subsequently used for Wards clustering (JMP13) in each dimension to generate dendrograms that robustly depict how samples or features cluster and should be ordered for a given choice of N clusters, which can then be overlayed aside the heatmap generated using the original Z scores. For two-way clustering one set of Z scores calculated from the same dimension (e.g. across samples for each gene) are independently subjected to consensus clustering to define the dendrograms and order for both features and samples, which are combined to generate a final heatmap.

**Chip-seq**

For Chip-seq experiments, PJ34-iICN1 cells were seeded into six T175 flasks at 6 million cells each and on the following day cells were treated with 1000 ng/ml doxycycline for 36 h to induce ICN1 expression before scraping and processing cells for Chip-seq according to our detailed published protocol (45). Briefly, the processed sample was divided into two equal parts. One part was incubated with a rabbit monoclonal antibody against cl-NOTCH1 (Cell Signaling, #4147) to immunoprecipitate ICN1 cross-linked to DNA, while the other part was treated with purified Rabbit IgG as a negative control for background signal subtraction. Following washes and reverse crosslinking, DNA was eluted, purified, quantitated and used to generate libraries for next generation sequencing performed on a HiSeq 3000 instrument. Raw reads were aligned to hg19 using Bowtie, 30 million reads were randomly sub-sampled, and peaks called with the Model-based Analysis for Chip-seq tool in Python.

**Additional Statistical Analyses.**

The log 2 transformation was used for all count data and the logit transformation for all percentage data before statistical testing. GraphPad Prism or JMP13 statistical software was used for most analyses. Two-sided t-tests were employed for comparisons involving only two groups, whereas experiments with multiple groups were analyzed using analysis of variance (ANOVA). For the latter, a post-hoc Tukey test was applied for pairwise comparisons, or Dunnett’s test was used when comparing groups against a control treatment. Single sample gene set enrichment scores (ssGSEA) were calculated through the BROAD Institute’s Gene Pattern public server at https://www.genepattern.org/ using published lists specific to individual cell types that we vetted through cross-correlation of gene expression across >.9,000 solid tumor samples from the TCGA.
